# A modular DNA toolkit for single-molecule biophysics

**DOI:** 10.64898/2026.10.09.757888

**Authors:** Jaco van der Torre, Roman Barth, Klaudia Majszczyk, Thejas Satheesh, Brian T. Analikwu, Clément Rouillon, Eva Bertosin, Benjamin D. Rowland, Cees Dekker

**Author notes:** Institute of Protein Design, Department of Biochemistry, 98195, Seattle, WA, USA. Bijvoet Centre for Biomolecular Research, Chemistry Department, University of Utrecht, Utrecht, 3584CH, The Netherlands (Eva Bertosin).

## Abstract

Long, customizable DNA molecules are an essential tool for in vitro single-molecule biophysics. Most studies still rely on the λ-phage genome, which has a fixed sequence, a non-uniform GC distribution, and limited options for modification. To overcome these limitations, we developed a modular DNA (ModDNA) toolkit that integrates in silico sequence design with cloning, to create long (up to 100 kb) DNA molecules with entirely user-defined sequences. We used a computational workflow with curated databases to design a motif-depleted sequence backbone, into which any desired sequence can be inserted at any position. Golden Gate assembly then builds these sequences into constructs, which we extend iteratively in defined increments up to 100 kb. We show precise placement of local features, e.g. a fluorophore, a bound protein, and a DNA-origami structure, and illustrate the construct’s utility in single-molecule visualization assays, including DNA loop extrusion by human condensin I. ModDNA offers a scalable, sequence-programmable alternative to λ-DNA and other restrictive substrates. More broadly, the motif-aware feature-free design connects single-molecule biophysics to emerging efforts in machine-guided regulatory-sequence design and synthetic-genome engineering, where controlled programmable sequence context is increasingly essential.

**Graphical abstract:** 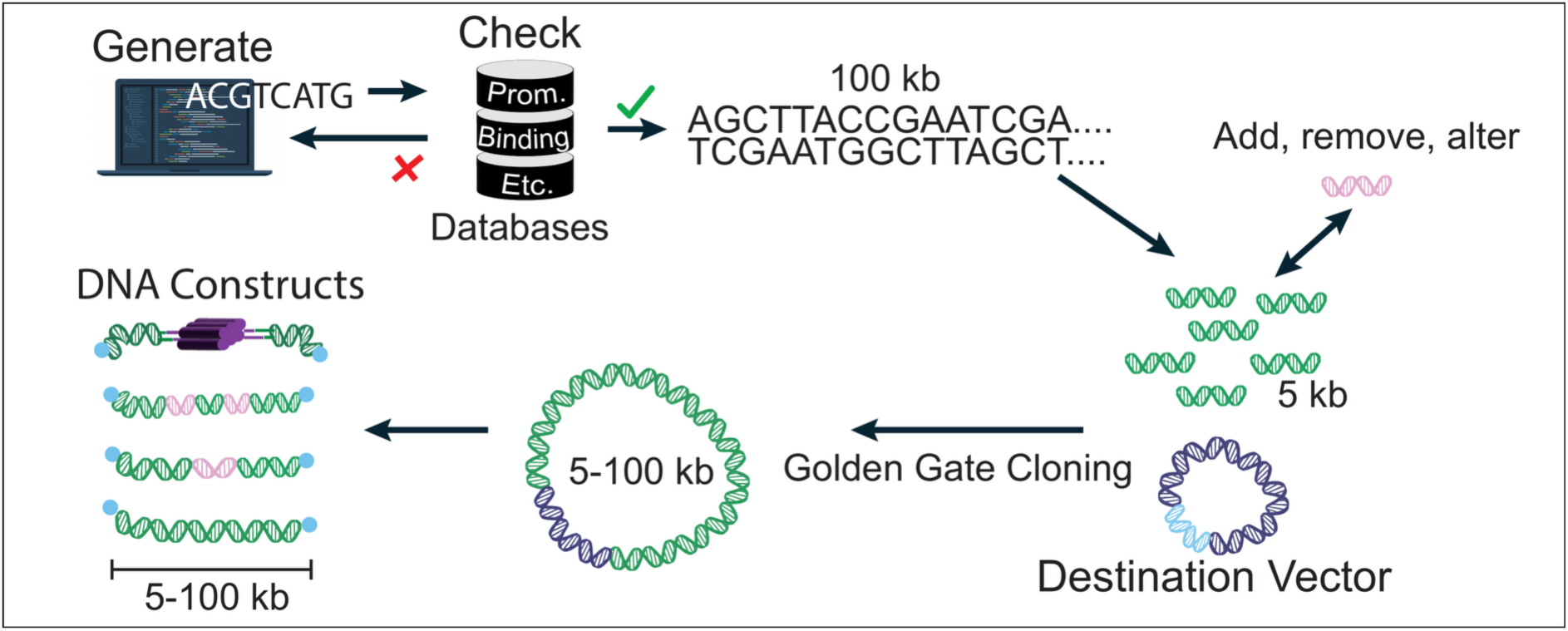

## I. Introduction

DNA sequence determines biological function via many mechanisms. While coding for proteins is the most familiar, the majority of the genome is non-coding, often exhibiting regulatory functions. Indeed, many non-coding DNA elements such as promoters, enhancers, insulators, telomeres, response elements, CpG islands, silencers, and other regulatory elements, contribute to the three-dimensional genome organization and control how genetic information is processed by the cell (1). The complexity of these factors hampers direct investigation of the molecular mechanisms.

For decades, biophysicists have developed *in vitro* single-molecule assays to study the isolated effects of individual elements on DNA. For example, the mechanical properties of DNA and DNA-binding proteins have been characterized using optical tweezers (2), magnetic tweezers (3), fluorescence imaging (4), and combinations thereof (5), facilitating a detailed investigation of how DNA’s physical properties, including sequence effects, influence its function. Single-molecule studies of DNA-protein interactions have significantly advanced our understanding of fundamental biological processes such as DNA replication (6), transcription (7), nucleosomes (8,9), RNA polymerases (10), transcription factors (11), and molecular motors such as structural maintenance of chromosomes (SMC) proteins (12,13). The development of an intercalation-based single-molecule fluorescence assay (14) enabled to study the role of DNA sequence in plectoneme formation (15) and dynamic DNA loop extrusion by SMC complexes such as cohesin and condensin (12,13,16,17). Alternatively, DNA curtains also visualize long DNA and bound proteins (16). All these experiments require well-defined DNA substrates of varying lengths.

Traditionally, the λ-phage genome (λ-DNA, with a length of 48,502 bases) has served as the standard DNA substrate for most studies involving long DNA molecules, due to its length and facile availability. Its fixed sequence and limited modularity do, however, restrict experimental design, particularly when specific sequences or functional elements are required. Various groups, including us, have demonstrated that specific DNA sequences can be engineered within λ-DNA or other long DNA substrates (18–20). However, manipulating these substrates is challenging and limited for the following reasons. (i) The viral capsid capacity imposes constraints on experimental design: the packaging mechanism of λ-phage permits only approximately 38-52 kb and exceeding 105% of the 48.5 kb genome rapidly decreases the yield and infectivity, thereby restricting the size of engineered constructs (21). (ii) Targeted editing is challenging as site-specific, modular insertion into native λ-DNA is technically complex, and traditional cloning often requires multistep ligations that have low yields for these long molecules. Researchers therefore employ recombination-based toolkits and engineered λ-phage platforms to achieve precise placement of features (22,23), but the constraint of a fixed genome sequence remains. We sought to radically overcome the limitations imposed by λ-DNA’s fixed sequence and aimed to develop customizable DNA scaffolds that have the potential to significantly expand the scope of *in vitro* single-molecule research by facilitating the investigation of sequence-specific phenomena such as nucleosome positioning and transcription factors.

The custom design and synthesis of long DNA fragments with particular sequences is nontrivial. As mentioned above, modification of λ-DNA and alternative methods (22) retain many fixed sequences and restrict the number of modifiable positions. Advances in molecular biology, including Gibson assembly and Golden Gate cloning, have however facilitated modular DNA assembly by enabling precise and efficient ligation of multiple fragments in a single reaction (24–26). So far these techniques have been used to yield some constructs of ∼10 kb length for single-molecule assays (14,27,28), but they did not provide the lengths necessary to investigate processes spanning tens to hundreds of kilobases. The synthetic biology field has developed multi-fragment assembly systems such as MoClo (Modular Cloning), Golden Braid, and MIDAS (Modular Idempotent DNA Assembly) (29–32), which utilize the Golden Gate approach to construct complex genetic assemblies from standardized building blocks. These strategies were primarily optimized for genome replacement or synthetic biology applications, and enabled the construction of even megabase-scale synthetic genomes (33,34). Despite their potential, these frameworks have not yet been adapted for generating long DNA substrates for single-molecule biophysics experiments.

Beyond cloning strategies for the construction of long DNA molecules, careful selection of the sequence content is critical for assay design. Many proteins recognize specific DNA motifs (35), so unintended sites can confound an experiment: cryptic promoters, pause sites, or terminators complicate transcription studies, and spurious protein-binding sites can interfere with many single-molecule assays. An immense amount of sequence information is available, as the sequence landscape and its influence on gene regulation and protein binding have been extensively characterized, resulting in curated databases that annotate regulatory elements for many organisms ((35–48); Materials & Methods).

Here, we present the design and validation of a modular DNA (‘ModDNA’) toolkit for generating DNA molecules with an entirely custom-designed sequence and tunable length from 5 to 100 kb. We first generate DNA sequences *in silico* that exclude any undesired features as listed in a long series of curated databases. These random sequences are thus strongly depleted of annotated regulatory and protein-binding motifs, providing a well-defined low-background context for single-molecule experiments (and other applications). Next, we devise and implement a Golden Gate-based cloning scheme that enables the efficient assembly of these sequences into constructs of up to 100 kb in length. Finally, we validate our approach by testing the resulting ModDNA molecules in various single-molecule assays, demonstrating the versatility and effectiveness of ModDNA for a range of biophysical applications.

## II. Materials and Methods

### Database collection

In order to identify any biologically relevant undesired features in DNA sequences, we collected various databases (see below) to mine for (i) sequences related to transcription such as promoters, (ii) enhancers, (iii) transcription termination sites (TTS); (iv) transcription factor binding sites; (v) DNAse hypersensitive sites (DHS); and (vi) those with predicted secondary structure (such as hairpins) as these can potentially interfere with replication of the designed sequences. We defined the latter as sequences with a low minimum free energy, because such sequences can readily fold into a secondary structure that may interfere or inhibit replication. All databases were downloaded on 2 July 2023. Since the methodology is described here in detail, users can apply it to additional or more recent databases of their choice with the code that is provided through Zenodo.

#### Promoters

Promoter sequences were compiled from multiple curated databases, enlisted below, to create a comprehensive library of experimentally validated and computationally predicted promoter regions across multiple species:

i. EPDnew Database. Eukaryotic promoter sequences were obtained from the Eukaryotic Promoter Database (EPDnew (36)) hosted at the Swiss Federal Institute of Technology (EPFL). Promoter annotations were downloaded via FTP from ccg.epfl.ch for 16 organisms: *Apis mellifera*, *Arabidopsis thaliana*, *Caenorhabditis elegans*, *Canis familiaris*, *Drosophila melanogaster*, *Danio rerio*, *Gallus gallus*, *Homo sapiens* (coding and non-coding), *Macaca mulatta*, *Mus musculus* (coding and non-coding), *Plasmodium falciparum*, *Rattus norvegicus*, *Saccharomyces cerevisiae*, and *Schizosaccharomyces pombe*. For each organism, the most recent genome assembly version was selected (last checked on 02-07-2023). Promoter coordinates were provided in BED format and mapped to their respective UCSC genome assembly identifiers.
ii. Genome Assembly Retrieval. Reference genome assemblies were downloaded on 02-07-2023 from the NCBI GenBank database (ncbi.nlm.nih.gov/genomes/) to extract promoter sequences from genomic coordinates. Assembly identifiers were obtained by cross-referencing UCSC genome codes with NCBI accession numbers via the assembly conversion table hosted at https://ccg.epfl.ch/mga/AssemblyTable.html. Genome sequences were downloaded in compressed FASTA format and parsed using BioPython’s SeqIO module. For each promoter annotation, sequences were extracted with 50 bp flanking regions on both sides of the annotated coordinates to capture potential regulatory elements at promoter boundaries. Sequences on the reverse strand were converted into its reverse-complement counterpart to maintain 5’ to 3’ orientation.
iii. Prokaryotic Promoter Databases. Bacterial promoter sequences were obtained from four specialized databases. RegulonDB (version 11.0), (37) provided experimentally validated promoter sequences for *Escherichia coli K-12*, including strand orientation and genomic coordinates. PromEC (39) provided additional *E. coli* promoter sequences in raw format without genomic coordinates. The Prokaryotic Promoter Database (PPD) (40) contributed promoter sequences from diverse bacterial species with associated metadata including organism taxonomy, genomic position, and strand information. PRODORIC (Prokaryotic Database of Gene Regulation) (41) supplied promoter sequences from multiple bacterial species with organism taxonomy and promoter names.

For data integration and quality control, all promoter sequences were compiled into a unified BED-format file containing the following fields: database source, organism, chromosome, promoter name, strand orientation (+/-), start coordinate, end coordinate, and DNA sequence. The final promoter library was chunked into overlapping fragments of 20 nucleotides with 10 bp steps to enable comprehensive pattern matching across all potential promoter subsequences during sequence analysis.

#### Enhancers

Enhancer annotations were obtained from EnhancerAtlas 2.0 (http://www.enhanceratlas.org/) (42), a manually curated database integrating enhancer predictions and experimental validations across multiple species. EnhancerAtlas 2.0 aggregates data from histone modification ChIP-seq, chromatin accessibility assays (DNase-seq, ATAC-seq), and transcription factor binding profiles to identify high-confidence enhancer elements. Enhancer coordinates were downloaded for nine model organisms spanning vertebrates, invertebrates, and fungi: *Homo sapiens* (hg19), *Mus musculus* (mm9), *Drosophila melanogaster* (dm3), *Caenorhabditis elegans* (ce10), *Danio rerio* (danRer10), *Rattus norvegicus* (rn5), *Saccharomyces cerevisiae* (sacCer3), *Gallus gallus* (galGal4), and *Sus scrofa* (susScr3). Data integration was performed as for promoters.

#### Terminators

Terminator sequences were compiled from multiple databases for prokaryotic terminators: (i) de Hoon Dataset: Intrinsic (rho-independent) bacterial terminators were obtained from (47) (Dataset S4). This dataset contains experimentally validated and computationally predicted transcription terminators from diverse bacterial species, compiled from literature sources and genome-wide analyses. (ii) WebGeSTerDB (48), which provided a database of stem-loop structures functioning as transcriptional terminators in prokaryotic genomes. Terminator sequences in WebGeSTerDB are represented with structural notation indicating four functional components: US (upstream stem, the 5’ arm of the inverted repeat), B (bulge or loop region), DS (downstream stem, the 3’ arm of the inverted repeat), and T (poly-U tail region). These components were concatenated to generate complete terminator sequences. Sequences were provided separately for the regular strand and complementary strand, ensuring comprehensive coverage of terminators on both DNA strands. WebGeSTerDB sequences contained ambiguity codes in parenthetical notation (e.g., A(C) indicating a position that can be adenine or cytosine). These were converted to standard regular expression format by removing parentheses while preserving the nucleotide alternatives, generating patterns compatible with sequence searching algorithms. All RNA sequences (containing uracil) were converted to DNA format.

#### Restriction and nicking sites and repetitive sequences

We also excluded the recognition sites of enzymes reserved for downstream manipulation, so that each acts only where deliberately placed: the Type IIS assembly enzymes BsaI (GGTCTC) and BsmBI (CGTCTC) necessary for Golden Gate assembly; the nicking enzyme Nb.BbvCI (CCTCAGC), for nick-and-ligate oligonucleotide labelling; a TypeII sticky end restriction enzyme NotI (GCGGCCGC); and a Type II blunt restriction enzyme PmeI (GTTTAAAC). Furthermore, repetitions of 4 or more equal nucleotides were excluded.

#### DNase I hypersensitive sites

DNase I hypersensitive sites (DHSs) annotations were compiled from publicly available datasets to identify regulatory regions that should be avoided in synthetic sequence design. DHS annotations were obtained from multiple genome-wide mapping studies across five genome assemblies: For human genome build hg38, the DHS Index and Vocabulary catalog (WM20190703 release) was retrieved from ENCODE (43). Mouse DHS annotations were obtained encompassing 197 biological samples mapped to the mm10 genome assembly (SI files of (44))(45). Plant DHS data were retrieved for *Arabidopsis thaliana* (TAIR10 assembly) and *Brachypodium distachyon* (Bd21 MIPS assembly) from https://plantdhs.org/ (46). Data integration was performed as for promoters.

### Nucleic Acid Folding Energy Prediction and Filtering

Beyond avoiding known regulatory sequences, synthetic DNA constructs must also minimize unwanted secondary structure formation that could interfere with replication or transcription processes. Secondary structures in both RNA (during transcription) and DNA (during replication) were predicted and used as exclusion criteria during sequence design.

Nucleic acid folding energies were calculated using the seqfold package v0.7.15 (49), which implements nearest-neighbor thermodynamic models for predicting the minimum free energy of secondary structures. The Gibbs free energy (ΔG) of folding was computed for both DNA and RNA forms of each sequence fragment. For DNA sequences, the free energy represents potential hairpin, stem-loop, or cruciform structures that could form during replication or in single-stranded regions. For RNA sequences, the free energy represents potential secondary structures in the transcribed mRNA that could affect translation efficiency or mRNA stability.

To establish biologically meaningful folding energy thresholds, a benchmarking analysis was performed on randomly generated sequences that had passed all database-based exclusion criteria. Random sequences of length equal to the chunk size (10-20 nucleotides, matching the design parameters) were generated from all possible nucleotide combinations. Each random sequence was validated to ensure it did not match any forbidden patterns from the regulatory element databases or restriction enzyme sites before being included in the benchmark set. For a benchmark set of 100,000 sequences, folding energies were calculated at multiple physiologically relevant temperatures (20°C and 30°C).

The distribution of folding energies was characterized for both DNA and RNA across all benchmark sequences. Folding energies were plotted as histograms on both logarithmic and linear scales to capture the full range of thermodynamic stability (Figure S1A-F). Sequences exhibited a wide range of folding energies, with most showing positive or near-zero ΔG values (indicating little to no stable structure), while a minority formed stable secondary structures with negative ΔG values. Based on these distributions, rejection thresholds were set at ΔG < -3 kcal/mol for both DNA and RNA structures. This threshold was selected to exclude sequences forming moderately stable or highly stable secondary structures while allowing sequences with minimal or transient structural propensity.

### De Novo Sequence Generation and in silico Assembly

#### Stochastic Sequence Construction Algorithm

Synthetic DNA sequences were generated using a constraint-based stochastic algorithm that iteratively builds sequences while enforcing multiple exclusion criteria. The algorithm operates on short sequence fragments (chunks) of 20 nucleotides, which are randomly generated and tested against all exclusion criteria before being concatenated to the growing sequence. Random nucleotide chunks were generated by sampling uniformly from the four-letter DNA alphabet {*A*, *C*, *G*, *T*}. Each candidate chunk was appended to a validation window consisting of the last N nucleotides of the existing sequence, where N is twice the maximum length of any forbidden motif (typically 100-200 nucleotides). This sliding window approach ensures that forbidden sequences are also detected if they span chunk boundaries. The concatenated validation sequence (previous window + new chunk) was tested against three criteria: (i) regex pattern matching for transcription factor binding sites and restriction enzymes, (ii) direct sequence comparison against promoter, enhancer, terminator, and DHS databases, and (iii) secondary structure formation potential for both DNA and RNA forms. For each fragment, the folding energy was calculated for both the DNA sequence and its RNA equivalent (with T→U substitution) at the design temperature (typically 20°C). If any fragment exhibited a folding energy below the rejection threshold for either DNA or RNA, the entire sequence chunk was flagged and excluded from further consideration.

#### Constraint Satisfaction and Backtracking

If a candidate chunk violated any exclusion criterion, a new random chunk was generated and tested, repeating until a valid chunk was found or a trial limit (500 attempts) was reached. When 500 consecutive trials failed to identify a valid chunk, the algorithm performed a backtracking operation, removing the last 2-7 chunks from the sequence (with the number scaled by the count of previous backsteps) to escape local sequence space constraints. This backtracking mechanism prevents the algorithm from becoming trapped in sequence contexts where no valid continuation exists within the allowable chunk size. To minimize internal sequence repetitiveness beyond that enforced by database comparisons, each newly accepted chunk was fragmented into overlapping subsequences of length (chunk size/2) and added to the exclusion list for the remainder of sequence generation. This dynamic expansion of forbidden sequences ensured that recently used sequence motifs were not reused, promoting sequence diversity and reducing the likelihood of spurious internal homology that could cause problems during DNA synthesis or cloning.

#### Sequence Fragmentation for Modular Assembly

Upon completion of the in silico sequence generation, the full-length construct (50-100 kb) was segmented into fragments of 5,000 nucleotides for subsequent practical DNA synthesis with Golden Gate assembly. Fragments were designed to be joined using Type IIS restriction enzyme-mediated assembly, which relies on 4-nucleotide overhangs to direct ordered fragment ligation. Fragment boundaries were positioned to yield pieces of uniform size (±2 nucleotides to account for restriction enzyme cleavage patterns), with the final 2 nucleotides at each fragment terminus reserved for overhang generation during restriction digestion. Fragment junctions were assigned unique 4-nucleotide overhangs, designated type C overhangs. The set was assembled for this work, and was checked with the NEBridge Ligase Fidelity Viewer (NEB, (50,51)). The 21 type C-overhangs (C0 to C20, given 5’ to 3’) are CGAG, AGCC, GTAT, TCAG, AAGT, TAAT, TCTA, TGTT, GCTG, TCGC, TACC, TTCT, TGAC, CGGT, TACG, GTGC, CCCT, GAGG, CTTT, TTTG and TTTC. 21 overhangs are required to join twenty 5 kb fragments, including the two terminal overhangs used for handle attachment. The same set supports assembly at every level of hierarchy: 5 kb donors into 10 to 20 kb intermediates, and intermediates into the complete 100 kb construct. The NEBridge evaluation used the BsaI and BsmBI-v2 cycling profiles. Since mis-ligation can only occur between overhangs present in the same reaction, we tested the most demanding case, a one-pot assembly containing all 21 overhangs; the staged assemblies each contain at most five. Predicted fidelity for the full set was 95 to 98%, and all 21 overhangs were classified as good Watson-Crick pairs. Each fragment was flanked by appropriate restriction enzyme recognition sites: BsmBI (CGTCTC) for fragment-level assembly and BsaI (GGTCTC) for higher-order assembly, with 10-nucleotide spacer sequences positioned 5’ to recognition sites to ensure efficient enzyme binding and cleavage.

After initial overhang assignment, each fragment junction (consisting of fragment N terminus, overhang sequence, and fragment N+1 start) was again validated against all exclusion criteria to detect forbidden sequences arising from overhang-fragment combinations. When violations were detected, targeted mutagenesis was performed on the fragment sequences immediately adjacent to the problematic overhang while keeping the overhang itself fixed. Mutations were introduced iteratively, starting with single-nucleotide substitutions and progressively expanding the mutation window (up to 10 nucleotides on each side of the overhang) until a valid sequence context was achieved or a trial limit (500 attempts) was exceeded. If local mutagenesis failed to resolve the conflict after 200 attempts, a fresh sequence fragment of equivalent length was generated *de novo* using the same constraint-based algorithm and substituted for the problematic region.

Following individual fragment validation, the complete assembled sequence (including all overhangs) was analyzed for forbidden motifs that might span multiple fragment junctions. Problematic sequences bridging fragment boundaries were identified and corrected through targeted mutagenesis of the nucleotides surrounding the relevant overhangs, with the constraint that overhang sequences themselves remained fixed to preserve assembly fidelity. Mutations were tested first in isolation (to ensure the mutated sequence itself was valid), then in a broader context (± 100 nucleotides), and finally across the complete construct to verify that corrections did not introduce new violations elsewhere in the sequence. After each successful correction, the modified sequence replaced the original, and the validation process continued until all inter-fragment conflicts were resolved.

The final validated sequence was output as a set of DNA fragments with complete flanking sequences for synthesis and assembly. Each fragment file contained the following adjacent sequence motifs: (1) a 10-nucleotide spacer, (2) the BsmBI or BsaI recognition sites, (3) a 1-nucleotide buffer, (4) appropriate overhang(s) based on assembly hierarchy, (5) the 5000-nucleotide core sequence, (6) the reverse complement of the 3’ overhang, (7) a 1-nucleotide buffer, (8) the reverse complement of the restriction site, and (9) a 10-nucleotide spacer. These fragments are compatible with commercial gene synthesis services and can be directly assembled using Golden Gate methods without additional processing using our ModDNA toolkit vector database.

### Analysis of given sequences

To analyze generated as well as pre-existing (e.g. naturally occurring) DNA constructs for the presence of regulatory elements, a reverse analysis pipeline was implemented. This pipeline searches input sequences against all compiled databases used during sequence generation, providing comprehensive identification of transcription factor binding sites (see below), promoters, enhancers, terminators, DNase hypersensitive sites, restriction enzyme recognition sites, and predicted secondary structures. Input sequences were searched against all pattern databases described above (CollecTF, promoters, enhancers, terminators, DHS, restriction sites, and repetitive sequences). For regex-based patterns (transcription factor binding motifs), patterns were compiled and applied using Python’s re.finditer() function. For direct sequence comparisons (promoter, enhancer, and terminator fragments), an optimized sliding window approach was employed: sequences were grouped by pattern length, and a set-based lookup (O(1) complexity) was performed at each window position rather than iterating through all patterns individually (which would be O(n) per position). Both forward and reverse complement strands were searched for all pattern types.

#### Transcription factor binding sites from JASPAR for sequence analysis

Transcription factor binding site motifs were obtained from the JASPAR 2026 database using the PyJaspar interface (35,38,52). We retrieved entries from the CORE collection for *Saccharomyces cerevisiae* (budding yeast). Each JASPAR entry contained a position frequency matrix (PFM) representing the nucleotide composition at each position of the binding motif. These count matrices were converted to frequency matrices by normalizing each column by its sum. To account for positions with zero counts across all nucleotides, column sums of zero were replaced with 1/4, and any remaining zero-sum columns after normalization were set to uniform frequencies (0.25 for each nucleotide), ensuring numerical stability in subsequent calculations. Frequency matrices were then transformed into Information Content Matrices (ICMs). First, zero frequencies were floored at 10^-3^. The information content at each position was calculated as *ICi* = 2 − *Hi*, where *H_i_* = − ∑*_b_*_∈(*A*,*C*,*G*,*T*)_ *f_b_*_,*i*_ log_2_3*f_b_*_,*i*_4 represents the Shannon entropy at position i, and *f_b_*_,*i*_ is the frequency of base b at position i. The factor of 2 represents the maximum possible information content for DNA sequences (log₂(4) = 2 bits). The final ICM was obtained by scaling each frequency by its position’s total information content: *ICM_b_*_,*i*_ = *f_b_*_,*i*_ · *IC_i_*. ICMs were converted to regular expression patterns for computational searching using a per-nucleotide information content threshold of 0.1 bits. At each position, individual nucleotides with IC values ≥ 0.1 were included in the pattern. Positions with a single nucleotide exceeding the threshold were represented as that specific base, while positions with multiple nucleotides above threshold were encoded using bracket notation (e.g., [A,C] for positions accepting adenine or cytosine). Positions where no nucleotide exceeded the threshold were represented as wildcards. To ensure pattern specificity, motifs with fewer than 5 informative positions (excluding wildcard characters) were excluded from the final pattern set.

This approach generated a comprehensive library of searchable patterns representing biologically relevant transcription factor binding sites while maintaining computational tractability for sequence analysis. A total of 294 regular expression patterns were generated in this way for *S. cerevisiae*.

#### PWM construction for sequence analysis

As an alternative to the discretized consensus-pattern search described above, the same set of JASPAR CORE motifs was also scanned using quantitative position-specific scoring matrices (PSSMs). Rather than collapsing each motif to a binary present/absent regular expression, this approach retains the full per-position, per-base weights and assigns each candidate window a continuous match score. The two analyses were run independently and their results are reported separately in Figure 2C-D and Figure S1H, as the two methods apply different criteria for calling a binding-site hit and are not directly comparable on a site-by-site basis. For each JASPAR motif, the position count matrix was converted to a position-specific scoring matrix (PSSM) of base-2 log-odds scores using Biopython’s PSSM engine. A uniform background (p = 0.25 for A/C/G/T) was used for the log-odds calculation. Pseudocounts were added before scoring via Bio.motifs.jaspar.calculate_pseudocounts. A separate PSSM was built for the reverse-complemented motif. For each motif the theoretical maximum and minimum attainable scores were recorded.

For sequence scanning, the query sequence was scanned with Biopython’s pssm.calculate(), which slides the PSSM over every window of the motif’s length and returns the summed per-position log-odds score (bits) for each window. Both strands were screened by scanning the same forward sequence with the forward PSSM and, separately, with the reverse-complement PSSM; reverse-strand hits are therefore reported directly in forward-sequence coordinates. Windows containing non-ACGT characters were excluded. A window was called a hit if its score met a relative threshold score ≥ min_score + f · (max_score − min_score), where f is a user-defined threshold (here 0.85) and min_score/max_score are that motif’s worst and best possible scores. This procedure normalizes the cutoff to each motif’s own dynamic range, because raw log-odds ranges generally differ between motifs of different length and information content.

### ModDNA plasmid cloning

Our cloning scheme utilized Golden Gate assembly, employing both BsaI and BsmBI type IIS restriction enzymes (NEB). The destination vectors were designed with a ‘sandwiched’ configuration (Figure 2F), placing BsaI and BsmBI sites on either side, both cutting at the same position. This arrangement enabled the generation of new donor vectors from all destination vectors. Golden Gate reactions were performed using either the BsaI-v2 or BsmBI-v2 Golden Gate Assembly Kits (NEB). Typical cycling conditions were 1 minute at 37°C (BsaI) or 42°C (BsmBI), alternating with 1 minute at 16°C, followed by a final incubation at 60°C for 5 minutes. Immediately after the 60°C step, reactions were transformed into NEB 5-alpha or NEB 10-beta chemically competent cells using standard protocols. For pBeloBac-based vectors, electrocompetent NEB 10-beta cells were used (NEB).

We initiated the cloning process by ordering our 100 kb sequence as 5 kb donor plasmids containing BsmBI sites, synthesized by GenScript in the pUC57mini-Amp backbone (Don2). Simultaneously, pUC57mini-Kan destination backbones were also obtained from GenScript. pUC57mini-Amp destination vectors were made from pUC57mini-Amp donor vectors and additional pUC57mini-Kan destination vectors were generated using site-directed mutagenesis (Q5, NEB) and KLD mix (NEB), or Gibson reactions using HiFi-mix (NEB), in combination with Genefragments (IDTDNA or Genscript). In total, 20 different pUC57mini-Amp based destination vectors were designed to facilitate the creation of alternative, short, and easily manipulated donor plasmids. The pUC57mini-Kan destination vectors were engineered to enable combination of multiple donor vectors, resulting in plasmids of 10–20 kb in size. For construction of the large plasmid, we modified the pBeloBac11 bacterial artificial chromosome plasmid, by combining dsDNA fragments (Genscript), PCR products, and Gibson assembly (HiFi, NEB) to yield a BsaI/BsmBI Golden Gate-compatible pBeloBac11 destination vector. Variants of these vectors were generated using site-directed mutagenesis (Q5, NEB) and KLD mix (NEB). Constructed plasmids were transformed in NEB 5-alpha or NEB 10-beta chemically competent cells using standard protocols, plated on LB agar plates with the appropriate antibiotic and grown overnight at 37°C. Subsequently, they were grown at 37°C overnight in LB and purified using a miniprep, midiprep, or Bac-midiprep kit from MN (Machery-Nagel); minipreps of cloned Bac-plasmids were purified without column purification. In short, 5 mL culture was centrifuged, followed by lysis, precipitation, and spin using MN-miniprep buffers and immediately followed by an Isopropanol-DNA-precipitation step including glycogen. After initial testing of plasmids using restriction enzymes and gel electrophoresis, all constructed plasmids were ultimately verified by whole-plasmid sequencing (Plasmidsaurus).

A summary of all plasmids constructed in this study is provided in Supplementary Table S1. Detailed information, including plasmid maps, sequences, and a complete list of primers used for cloning, is available in the Snapgene files of all plasmids that accompany this work (Zenodo). Plasmid features, assembly strategies, and relevant genetic elements are described in the table legend.

### DNA preparation for single-molecule assays

λ-DNA for single-molecule experiments was prepared as described previously (53). λ-DNA containing a biotin on both ends, was made by hybridizing and ligating short oligonucleotides containing 5ʹ phosphate and 3ʹ Biotin on the single-stranded DNA ends of λ-DNA (NEB, N3011S). For this we used oligonucleotides, JT41 (P-GGGCGGCGACCT-Bio) and JT42 (P-AGGTCGCCGCCC-Bio) (IDT). Taq DNA ligase (NEB, M0208L) was used to ligate the oligonucleotide on λ-DNA, using 10-fold molar access of oligonucleotide over λ-DNA. The mixture was incubated for 10ʹ at 65°C and then 1hour at 50°C in Taq DNA ligase buffer. The biotin-λ-DNA was then cleaned up from free oligonucleotides and enzymes using an AKTA start system (Cytiva), with an S500HR gel filtration column (Cytiva), run with TE + 150mM NaCl buffer at 0.5mL/min.

Plasmids were processed in a similar process as how λ-DNA constructs are made, with some key differences. Plasmid DNA was first treated with RecBCD for 30minutes at 37°C, followed by heat-inactivation 60°C for 10minutes. This step removes damaged plasmid DNA as well as *E. Coli* genomic DNA, which is a common contaminant when isolating large BAC plasmids. The ModDNA-BAC plasmids were cut with BsaI-v2 or BsmBI-v2 (NEB), heat inactivated 65°C for 10minutes and subsequently a ligation reaction was setup with the correct handles. Handles were duplex oligonucleotide carrying a 5’ phosphate and a 3’ biotin, with the complementary strand either unmodified or carrying an additional 5’ biotin (IDT). This gives one construct end with a single biotin and the other end with two biotins, which makes unconstrained biotinylated constructs. An approximate 10-to 100-fold excess of oligonucleotides was used. This was incubated overnight at 16°C. Alternatively for the construct with fluorescent handles, the handles were made by a PCR of 250nt, where during the PCR we had 1/5 Atto643-x-dUTP (Jenabioscience). These were digested with BsaI-v2, purified using a PCR-cleanup kit from NEB and ligated instead of the duplex-oligonucleotides. After ligation using T4 DNA Ligase (NEB), 10x excess, purification was performed using an S500HR gel filtration column (Cytiva). The procedure was conducted with an AKTA-start at room temperature, and separation capacity was tested with a DNA marker to ensure efficient removal of free handles (Figure S2).

### Internal labeling of construct with Biotin or AF647 for single-molecule assays

To introduce an internal biotin, internal Atto647N, or a DBCO-AF647 (Jena Bioscience) fluorophore into the DNA construct, we used the nicking-enzyme-based internal-labeling method of Luzzietti et al. (54). A 40kb pBeloBac11-ModDNA plasmid (plasmid Mod324) carried 5 BbvCI sites engineered in its centre, allowing the plasmid to be nicked whereupon a longer oligonucleotide bearing any modification could be ligated in. Three oligonucleotides were used for this: JT769(/5phos/TGAGGtttcGCTGAGGacgcGC/iBiodT/GAGGatcagagGCTGAGGatggagtGC), JT770(/5phos/TGAGGtttcGCTGAGGacgcGC/iAzideN/GAGGatcagagGCTGAGGatggagtGC) and JT771 (/5phos/TGAGGtttcGCTGAGGacgcGCTGAGGatcagagGCTGAGGatggagtGC/3ATTO647N) (IDT). JT770 was first reacted with DBCO-AF647 (Jena Bioscience) by copper-free click chemistry for 1 hour at room temperature, before being hybridized to the construct. Plasmid Mod324 was first treated with RecBCD for 30minutes 37°C, followed by heat-inactivation 60°C for 10minutes. It was then digested with BsmBI-v2 (NEB) for 2 h at 55°C. Nb.BbvCI was subsequently added to the same reaction and incubated for 30minutes at 37°C, followed by heat inactivation for 20 minutes at 80°C. Oligonucleotides JT769, JT770(+AF647) or JT771 were then added at a 100:1 molar ratio relative to the DNA construct and hybridized for 3 h at 40°C. Biotinylated handles were then ligated on and the construct was processed further as described above for DNA preparation for single-molecule assay.

### DNA origami nanostructure

One example of an included feature was a folded DNA origami nanostructure, an 80 nm-long 3x2-helix bundle designed using a square lattice using caDNAno (Figure S4A, (55)). Twist was adjusted by inserting a skip every 32 nucleotides in each helix. Two oligonucleotides at the interfaces (helices 0 and 1) were phosphorylated at the 5’ end, while five oligonucleotides in the structure core were modified with fluorescent dyes. To assemble such a small DNA origami, we identified two unique restriction sites within the p7560 scaffold (tillibit nanosystems GmbH), namely MscI and HincII, located 1426bp apart. We hybridized oligos (MscI: 5’ -GGG ACA TTC TGG CCA ACA GAG ATA-3’; HincII 5’ -TGC TGA ATT GTC AAC CTT ATG ACA-3’) to the single-stranded p7560 scaffold at a 5x excess. Restriction was carried out for 1 h at 37°C, followed by heat inactivation at 60°C for 20 min. The 1426 nt scaffold fragment was separated from the remaining 6.1 kb fragment on a 1% agarose gel and purified by gel extraction (NucleoSpin Gel and PCR Clean-up, Macherey-Nagel).

The folding reaction contained 40 nM scaffold (1426 nucleotide long), 200 nM oligonucleotides (IDT Integrated DNA Technologies), folding buffer (1 mM EDTA, 5 mM NaCl, 10 mM Tris-HCl (pH 7.5)), and 20 mM MgCl2. The folding reaction was heated at 65°C for 15 min and annealed from 60°C to 20°C (1°C per hour) in a thermocycler (T100, Biorad). The reaction was left in the fridge before and after purification. Excess oligonucleotides were removed via ultrafiltration (56) using Amicon Ultra 0.5 mL Ultracel filters 50K, using 1xFoB5 buffer (10 mM Tris-HCl, 1 mM EDTA, 5 mM NaCl, and 5 mM MgCl2). All the centrifugation steps were performed at RT, at 10k g for 3 min. First, the filter was washed with 500 µL of 1xFoB5. The permeate was discarded, and 50 to 100 µL of sample was pipetted into the filter. Buffer was used to top up the solution to 500 µL and centrifuged again. After discarding the permeate, 450 µL of 1xFoB5 was added and centrifuged. This step was repeated 5 times in total before retrieving the sample, placing the filter inset upside down in a new tube and centrifuging for 5 min. Samples were electrophoresed in gels containing 2% agarose, 0.5xTBE (Tris-borate-EDTA), and 5 mM MgCl_2_ for 1.5 hours at 90 V bias voltage. The gels were imaged in a gel imager (BioRad). Image processing was performed with FIJI (57). For TEM imaging, a carbon Formvar grid (Electron Microscopy Sciences) was glow-discharged (1 min, 45 mA) before use. 4 µL sample (concentration between 20/30 nM) was incubated onto the grid for 2 min. The excess sample was blotted with a filter paper. A 4 µL droplet of stain solution (2% uranyl acetate) was pipetted onto the grid and immediately blotted. Another 4 µL droplet of stain was pipetted onto the grid, incubated for 30 s, and blotted away. The grid was left to dry in air for minimum 2 min before imaging on a Jeol FSC3200 microscope. Class averages were computed using Relion 3.0 (58) (Figure S4 B-C).

To incorporate the DNA origami into the long DNA, two 20kb ModDNA constructs each carrying a biotin handle, on one side, were made, purified and subsequently ligated in a 2:1 ratio to the DNA-origami construct. For this, plasmid Mod303 was digested with BsmBI-v2, 55°C and heat-inactivated at 65°C for 10 minutes. Duplex oligonucleotides carrying a biotin were ligated to one side. We then purified DNA using an S500HR gel filtration column (Cytiva), in TE supplemented with 150mM NaCl, using an AKTA-start (Cytiva) at room temperature.

Loop extrusion on these constructs was performed as prescribed below, and the blocking efficiency of the DNA origami was determined in the same way as described in Analikwu et al. (18).

### Purification of yeast condensin, LacI, and human Condensin I

*Saccharomyces cerevisiae* condensin and *Escherichia coli* LacI were purified and fluorescently labelled as described by Analikwu et al.(18). Human condensin I was purified from HAP1 cells (59), in which CAP-H was endogenously tagged at the C-terminus with SNAP-3xFLAG using CRISPR-Cas9 technology. Oligonucleotides encoding guide RNAs targeting the C-terminus of CAP-H (5’ CAAGGAGATTGAGTTCACTA-3’) were cloned into pX330 and co-transfected with a repair template in pUC19 using FuGENE 6 (Promega). To select clones, cells were treated with 2 μg/ml puromycin for 2 days before picking colonies. Insertion of the tag into the correct locus was validated by Sanger sequencing of the isolated genomic DNA. Cells were cultured in roller bottles (Greiner, #681670) in Iscove’s modified Dulbecco’s medium (Invitrogen) supplemented with 10% fetal calf serum (FCS; Clontech) and 1% penicillin-streptomycin (Invitrogen). Cells were arrested in mitosis using nocodazole (100 ng/mL) for 18h before harvest. Next, 3-4mL pellet was lysed in lysis buffer (20 mM HEPES pH 7.5, 500 mM NaCl, 1 mM EDTA, 5% glycerol, 0.1% Tween-20, and 1 mM TCEP), supplemented with protease inhibitor (Roche). The cells were lysed by Dounce homogenization, followed by sonication. Lysate was then clarified by centrifugation at 56,000 x g for 40 minutes. The clarified lysate was incubated with anti-FLAG M2 agarose resin for 1 hour at 4° C while rotating. The sample was then loaded onto a gravity-flow column and the flowthrough was collected. Condensin I was labelled on beads with 5 µM SNAP-Surface Alexa Fluor 647 ligand (NEB) for 1 hour at 4° C Beads were washed with 40 column volumes of lysis buffer and 10 column volumes of wash buffer (20 mM HEPES pH 7.5, 150 mM NaCl, 1 mM EDTA, 5% glycerol, and 1 mM TCEP). Condensin I was eluted using 5 column volumes of wash buffer supplemented with FLAG peptide (0.5 mg/mL). The eluate was concentrated (AMICON MWCO 50 kDa), snap-frozen in liquid nitrogen and stored at -80°C. Concentration was determined via Coomassie-stained SDS-PAGE using a standard of known concentration. The integrity of the purified condensin I complex is shown on a Coomassie-stained gel (Figure S3A), and its composition was assessed by mass photometry (Figure S3B), which showed a main species of 666 kDa, consistent with the five-subunit complex.

### Preparation of LacI-LacO construct for single-molecule experiments

To label LacO sites prior to the loop extrusion assay, the DNA construct prepared for single-molecule assays (Don4b-C0-2x4LacO-C10, ∼5 pM) was incubated with 6 nM LacI-SNAP-AF647 for 10-15 min at room temperature in LacI buffer (100 mM KGlu, 20 mM Tris-HCl pH 7.5, 2.5 mM MgCl_2_, 0.25 mg/ml BSA, 1 mM TCEP). After incubation, SYBR Gold was added to 10 nM (final), the DNA was introduced in the flow channel, and the loop extrusion assay was performed as described below, with yeast condensin.

### Localization of LacO and Oligonucleotide-Atto647N Labels on DNA

The positions of LacO/LacI arrays and the Atto647N-oligonucleotide JT771 were verified using microscopy. DNA constructs were introduced into the flow channel following standard procedures. Unbound DNA and protein were subsequently washed out, and flow was halted. Recordings of DNA (imaged with 10 nM SYBR Gold using a 488 nm laser) and either LacI-SNAP-AF647 or Atto647N (using a 639 nm laser) were acquired. Single DNA molecules were cropped and background subtracted, then kymographs were generated. The positions of LacI and Atto647N-oligonucleotide were tracked from these kymographs. Each track produced a static but noisy position, which was averaged to obtain a per-track position. The locations of the LacO array or Atto647N were then inferred by fitting a double Gaussian probability distribution, and the mean and standard deviation were calculated.

### Loop extrusion assays

Loop extrusion assays were carried out in flow channels as previously reported (12). Briefly, flow channels were prepared by cleaning, functionalizing, and passivating 76.2x25.4 mm (3x2 in) slides and 25x50 mm coverslips. Slides contained typically 10 holes on either side that serve as inlets and outlets to flow channels. The slide cleaning procedure is comprised of successive rounds of rinsing and sonication in acetone, Millipore water, 1 M KOH, Millipore water, followed by acid piranha treatment. The functionalization procedure is done by aminosilanization with 5% APTMS (Sigma Aldrich), 10% glacial acetic acid (Boom), in methanol (Boom) for 20min, followed by 4 thorough rinses in methanol before baking at 70°C for 30min or until no visible methanol film is present. Passivation was done by PEGylating in 800 μg/ml mPEG5000-SVA and 20 μg/ml Biotin-PEG5000-SVA in 100 mM NaHCO_3_ and 550 mM K2SO4 buffer. A subsequent round of PEGylation was done with 300 mM M(PEG)_4_-NHS in the same buffer.

Flow cells were assembled by sandwiching a double-sided tape between the PEGylated slide and coverslip. The double-sided tape serves as the wall edges between individual channels. Tubing was attached to the outlet holes and connected to a syringe pump for constant flow inside the channels. Channels were incubated with 0.01 mg/ml streptavidin (NEB) and 0.5 mg/ml UltraPure BSA (Invitrogen) in T20 buffer (10 mM Tris pH 8, 20 mM NaCl) for at least 15min. The flow channels were washed with T20 and DNA with origami roadblocks was introduced in the flow cells at 3-5 μl/min flow rate in loop extrusion buffer (50 mM KGlu, 20 mM Tris-HCl pH 7.5, 2.5 mM MgCl_2_, 0.25 mg/ml BSA, 1 mM DTT, 10 nM SYBR Gold). Purified budding yeast condensin was introduced at 100-500 pM, condensin I was introduced at 200-400 pM in loop extrusion buffer supplemented with 1 mM ATP, 2 mM Trolox (UV activated to >10%), 10 nM catalase, 18.75 nM glucose oxidase. Buffer flow was stopped after the introduction of condensin for the observation of loop extrusion. Imaging was done on a HILO microscope, with a red (639 nm, 20 mW) and a blue (488 nm, 0.5 mW) laser, in alternating light excitation mode.

### Data analysis of loop extrusion

Loop extrusion analysis was carried out using a custom-built Python software as previously reported (18). Initially, we generated kymographs of the selected DNA molecule and tracked the peak intensities based on the DNA signal. The amount of DNA in the loop is obtained by multiplying the fraction of peak intensity with the length of the DNA (L) used (50 kb or 100 kb). In other words,

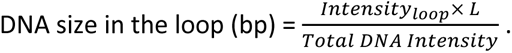

The loop tracking records for each molecule were loaded into another Python script, where we did further analysis. The changes in DNA size as a function of time (loop size traces) were smoothed with a Savitzky-Golay filter (21-frame window, first-order polynomial) to reduce noise. For each track, we chose a single, first, loop growth phase which was a period of processive loop extrusion. This was done using either manual selection or automatic detection. Manual selections were made by clicking at the start and end points of a growth phase. For automatic detection, we scanned the loop size trace with a sliding linear regression window (20 frames) and classified frames as growing wherever the local slope exceeds a certain threshold. This growth mask was cleaned and used for further downstream analysis. It is to be noted that using either manual or automatic selection did not yield significantly different values. From these growth phases, the steepest slope was taken as the loop extrusion rate. The maximum loop size was obtained from the largest loop size sustained over at least five frames in the whole track. Each phase was additionally tagged with the DNA molecule’s initial relative extension (fraction of contour length) based on their end-to-end distances. This was then used to generate all the plots.

### Worm-like-chain tension model

DNA was held between two fixed points such that the end-to-end distance was constant throughout the simulation. Initially, the DNA was held at a fractional extension *FCL* (fraction of contour length)

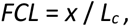

where *x* is the end-to-end distance and *L_c_* is the contour length of the DNA. At the start, the relative extension *Z_0_* was

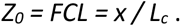

As condensin extrudes a loop of size *L_loop_*, the DNA outside the loop was under a growing tension. The contour length of the DNA outside the loop decreased to (*L_c_* − *L_loop_*), while *x* remained the same. The relative extension therefore grew to

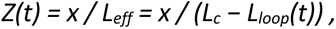

where *L_eff_* is the effective contour length of DNA under tension. *Z*(*t*) was then plugged into the extension term of the Marko-Siggia interpolation formula for the WLC model to obtain the force,

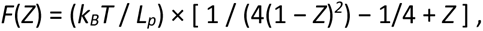

where *L_p_*is the persistence length and *k_B_T* thermal energy.

In the model, 2000 evenly spaced loop-size values were generated from 0 till close to the geometric limit where *L_eff_* = *x* (i.e. *Z* → 1). The relative extension was evaluated at each point, and the corresponding force was calculated from *F*(*Z*).

## III. Results

### Designing DNA from the bottom up

Prior to constructing the ModDNA molecules, we first extensively identified sequence features in silico to exclude from the design for the synthetic sequence (Materials and Methods, Figure 1 and S1). We engineered the sequence to eliminate as many sequence elements as possible, creating a “blank slate” sequence suitable for diverse applications. Unlike tools such as GeneDesigner, which prioritize codon optimization or expression (60), our design strategy focuses on removing as many protein-binding sites, regulatory elements, and stable secondary structures as possible. The resulting sequence is designed to be functionally inert, making it a clean building block for single-molecule experiments (and other applications).

**Figure 1.**
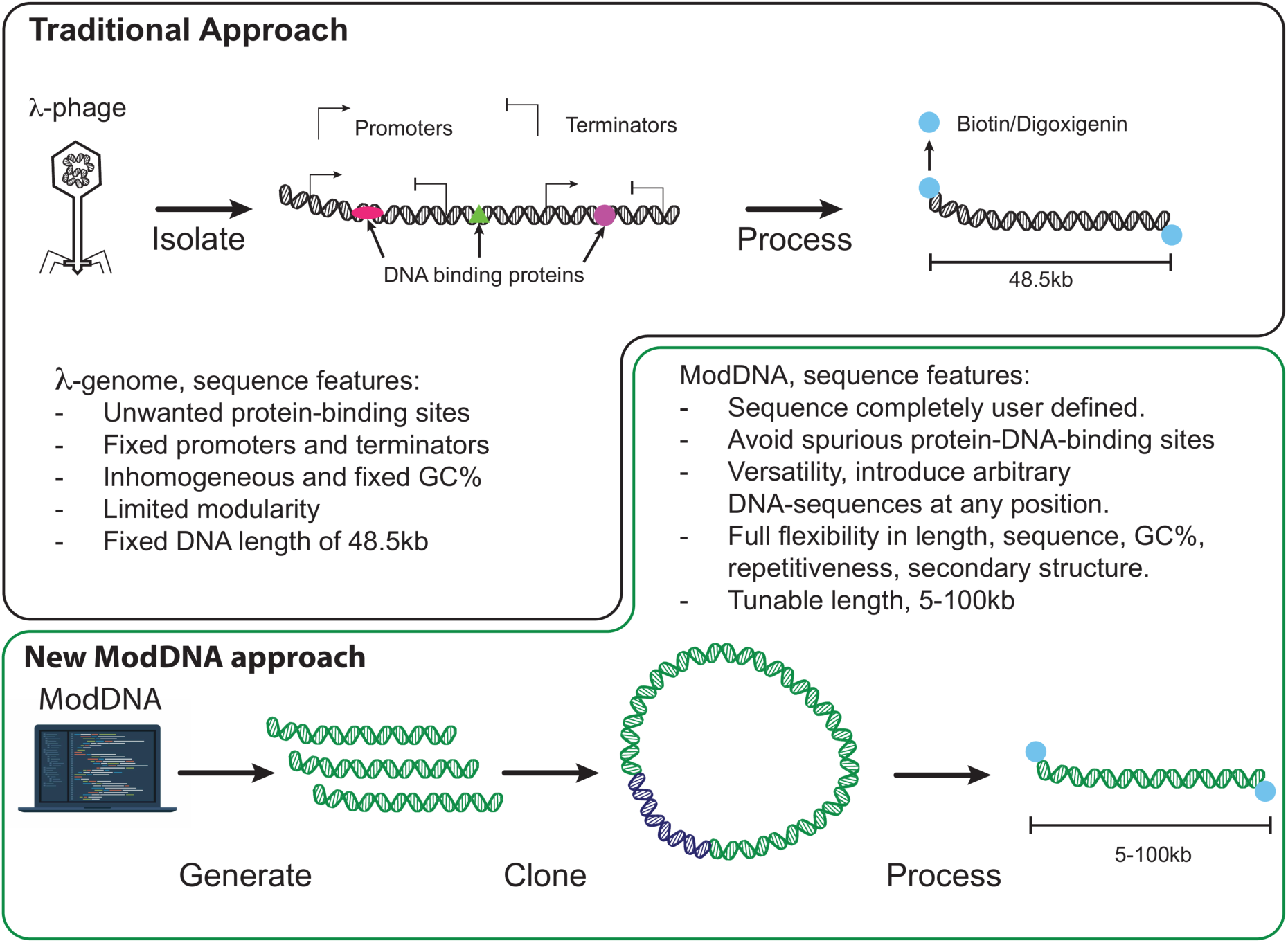
Design principles and modularity of the ModDNA system versus λ-DNA. Comparison of the traditional λ-DNA and the new ModDNA approaches in terms of DNA sequence flexibility and design constraints. λ-DNA contains fixed promoters, terminators, cryptic promoters, an anomalous GC content and distribution, and multiple protein-binding motifs, and it has a fixed length of 48.5 kb. ModDNA enables fully user-defined sequence composition, including adjustable GC content, (non)repetitiveness, and secondary structure, while avoiding spurious protein-binding sites. ModDNA constructs can be generated at arbitrary lengths between 5–100 kb.

Specifically, we used a constraint-based stochastic algorithm to generate a random synthetic 100 kb DNA sequence that explicitly excludes transcription factor binding sites, promoters, enhancers, transcription terminators, DNase hypersensitive sites, curated restriction enzyme recognition sites, and sequences predicted to form stable secondary structures (ΔG < -3 kcal/mol for DNA or RNA; Materials and Methods; Figure 2A). The algorithm iteratively generated random 20-nucleotide segments, validating each against these exclusion criteria using a sliding-window analysis, and employing backtracking when the local sequence space became constrained. To facilitate robust construction, the final sequence was divided into 5,000 bp fragments compatible with Golden Gate assembly. These segments were joined by 4-nucleotide overhangs selected for high predicted ligation fidelity (Materials and Methods). We validated and corrected the overhang junctions through targeted mutagenesis to ensure that no forbidden motifs spanned the overhang boundaries (Materials & Methods; Figure 2B). This process yielded a 100 kb sequence devoid of the many functional and cryptic DNA elements present in native DNA such as λ-phage (Figure 1).

**Figure 2.**
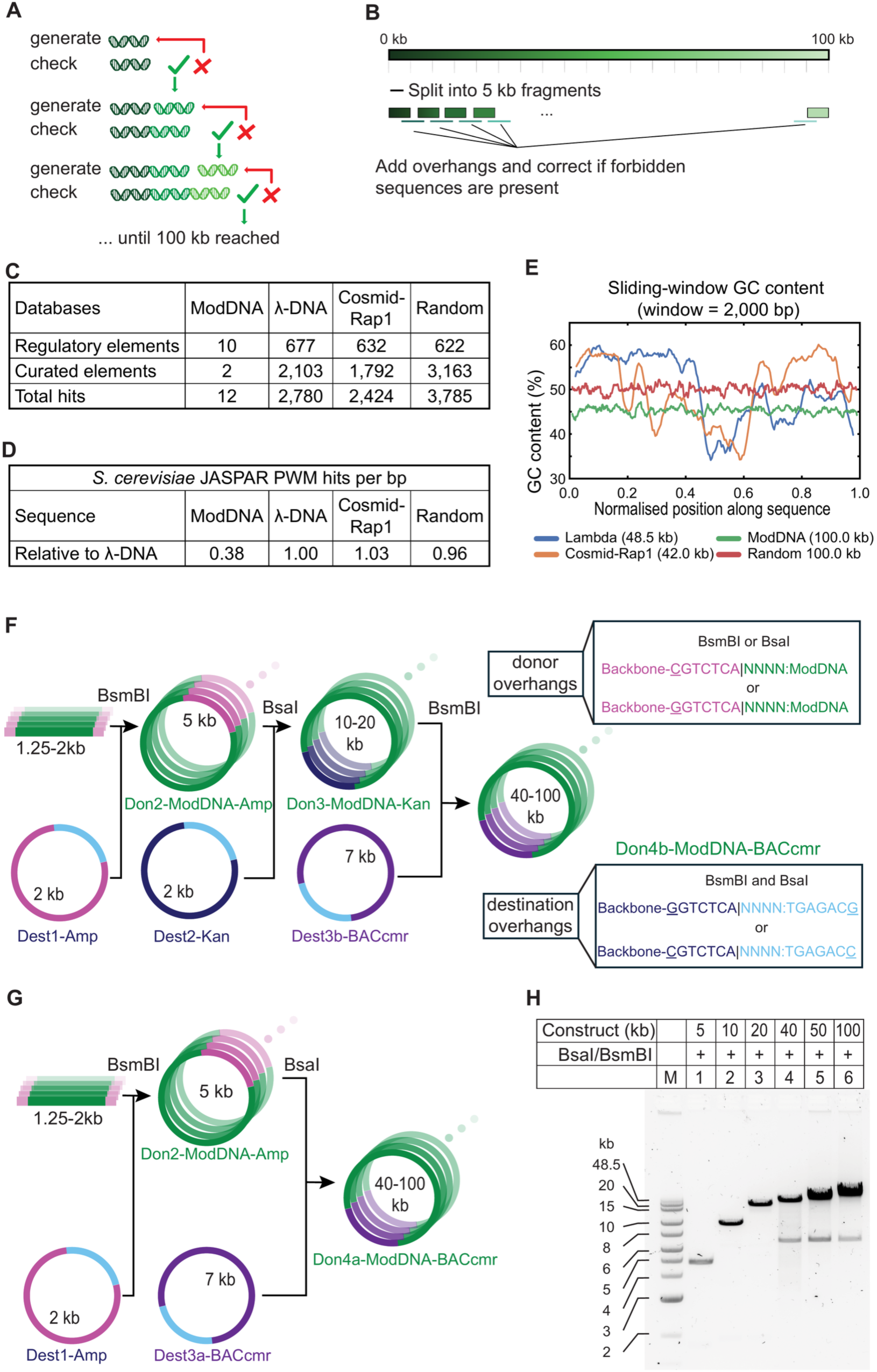
Assembly workflow of ModDNA constructs. (A) Stepwise computational workflow to generate a ModDNA sequence. DNA fragments of 20 bp are generated, sequence-verified for features, and iteratively combined until the target construct length (up to 100 kb) is reached. Newly created regions are screened against all databases used to generate the original 100kb sequence. (B) The resulting 100-kb sequence is segmented into 5-kb fragments, each appended with curated 4-nt overhangs for Golden Gate cloning. (C) Number of hits against the regulatory and curated element databases in λ-DNA, a Cosmid-i95rap1-DNA (18), ModDNA, and a random 100 kb sequence of 50% GC content, given as the total number of hits (not normalised to sequence length). (D) Density of predicted transcription-factor binding sites in the same four sequences. Each sequence was scanned with position weight matrices (PWM) from the JASPAR CORE collection (2026 release) for Saccharomyces cerevisiae at a relative-score threshold of 0.85 (Materials and Methods); hit counts were normalised to sequence length and are expressed relative to λ-DNA, which is set to 1. (E) Sliding-window GC content (2,000 bp window) along each sequence, plotted against normalised position. λ-DNA and Cosmid-Rap1 vary markedly along their length, whereas ModDNA holds a stable profile close to its global mean of ∼45% GC. (F) Hierarchical cloning schematic showing the architecture of the Type IIS restriction sites (BsaI, BsmBI) and the staged assembly route. Multiple donor vectors are combined with a single destination vector to generate a new donor vector. All donor vectors are suitable for single-molecule experiments. (G) Cloning schematic for the direct route, in which the 10-20 kb intermediate step is skipped and 5 kb donors are assembled directly into 40-100 kb constructs. This strategy was used to make three different ModDNA constructs, but it is less efficient than the hierarchical assembly strategy. (H) Agarose gel electrophoresis of assembled constructs ranging from 5 to 100 kb after digestion with either BsaI or BsmBI. The data demonstrate distinct, correctly sized products across the entire length range. Smaller bands correspond to vector backbones.

The suitability of ModDNA as a general substrate for single-molecule experiments was first assessed in silico by quantifying its motif burden, transcription factor binding sites, GC content distribution, and by comparing these properties with those of some DNA substrates in routine single-molecule use, λ-DNA, Cosmid-Rap1 (18), and with a randomly generated 100 kb sequence of 50% GC content (Materials and Methods). ModDNA was, as described in Materials and Methods, designed by iteratively depleting motifs compiled from a range of datasets, including promoters, enhancers, terminators, DNase I–hypersensitive sites, curated restriction sites, homopolymer runs, and transcription-factor motifs from CollecTF.

In each of the non-depleted substrates these motif classes were found to be present thousands of times per construct, whereas the corresponding counts in ModDNA were strongly reduced (Figure 2C and S1H). For example, 2780 hits were counted in the 48.5 kb λ-DNA (about 5,700 per 100 kb) versus 12 hits for the 100 kb ModDNA. As an independent test, using motif definitions that were not targeted during design, each sequence was scanned with position weight matrices from the JASPAR CORE collection (2026 release), scoring *S. cerevisiae* transcription-factor binding sites at a relative-score threshold of 0.85. Hit counts were normalized to sequence length and expressed relative to λ-DNA. The results show that ModDNA retained only 38% of the λ-DNA binding-sites, a 62% reduction compared to λ-DNA (Figure 2D). By contrast, Cosmid-Rap1 and the random 100 kb sequence were indistinguishable from λ-DNA (103% and 96%, respectively), showing that this depletion is specific to ModDNA’s design and not a generic consequence of base composition. GC content followed the same pattern. In sliding-window analysis, λ-DNA and Cosmid-Rap1 both vary markedly along their length, with the local GC content spaning roughly 31–65%, with correspondingly large variation in local curvature. ModDNA instead displayed a stable profile close to its global mean (∼45% GC) along its entire length (Figure 2E), giving it uniform, and therefore more predictable mechanical and thermodynamic behaviour for single-molecule measurements.

### Golden Gate cloning enables 100 kbp-long custom-designed DNA

Next we developed a cloning strategy to synthesize the ModDNA 100 kb construct using Golden Gate cloning. By exploiting Type IIS restriction enzymes, such as BsmBI and BsaI, destination vectors were engineered with sandwiched BsmBI/BsaI recognition sites that generate the same 4-nucleotide overhangs, enabling precise and seamless ligation of DNA fragments (61). This modular design allows the direct reuse of assembled plasmids as donor vectors, thereby facilitating iterative and theoretically unlimited rounds of cloning while maintaining sequence fidelity (Figure 2F and G). The deliberate incorporation of 4-nucleotide overhangs during in silico design grants fine control over the assembly junctions, eliminating unwanted scar sequences and ensuring feature integrity within the stitched regions. The sequence design enforces a specific orientation, thereby preventing the introduction of unintended genetic features that could result from random fragment rearrangement. The overhang set was assessed computationally using the NEBridge Ligase Fidelity viewer (NEB (50,51)), which predicted an assembly fidelity of 95 to 98% for a single-tube reaction containing all 21 overhangs and effectively complete fidelity for staged assemblies used here, with all 21 overhangs classified as good Watson-Crick pairs.

To generate single-molecule constructs with the ModDNA toolkit, we engineered a suite of destination and donor vectors covering a wide range of assembly sizes. Three plasmid backbones were used: pUC57mini-Amp (Dest1 and Don2, GenScript), pUC57mini-Kan (Dest2 and Don3, GenScript), and a derivative of the bacterial artificial chromosome (BAC) backbone pBeloBac11gg (Dest3a, Dest3b, Don4a and Don4b), rendered Golden Gate compatible by removing all endogenous BsaI and BsmBI sites from the backbone (Materials and Method). The high-copy pUC57mini vectors covered the 5-20 kb ModDNA range and provided the starting material: twenty pUC57mini-Amp-ModDNA donor plasmids (Don2, GenScript) each carrying one 5 kb fragment of the 100 kb ModDNA sequence. Above 20 kb, the single-copy pBeloBac11gg destination vectors took over, giving access to constructs carrying 40-100 kb of ModDNA sequence (Figure 2F and G). Two routes to these large constructs were built into the design: direct assembly from the 5 kb pUC57mini-Amp-ModDNA donors (Don2) in a single reaction (Figure 2G), or a hierarchical route in which 10-20 kb intermediates were first assembled in pUC57mini-Kan (Dest2) and subsequently combined in pBeloBac11gg (Figure 2F). Assembly endpoints realized so far were 10, 20, 40, 50, and 100 kb (Materials and Methods, Figure 2F and G, Table S1). The fidelity of each assembly step was verified by whole-plasmid nanopore sequencing (Plasmidsaurus) and by gel electrophoresis (Figure 2H).

For the longest constructs, assembly efficiency benefited from the hierarchical route. Passing through 10-20 kb intermediates added a cloning step, but each individual step was more robust: the proportion of correct clones rose, and the screening burden fell accordingly. Cloning of the larger constructs was sensitive to procedural detail, whereas smaller assemblies proved robust; when the hierarchical scheme was followed and those details were respected, screening four colonies was typically sufficient to obtain a correct clone, although individual steps occasionally had to be repeated (Materials and Methods). The hierarchical route is not required, however, and the one-pot assembly was used successfully for several constructs (Table S1). Nor did the 5 kb donors themselves need to be ordered: they could equally be produced from two to three synthetic dsDNA fragments and assembled into pUC57mini-Amp (Dest1), which markedly reduced the cost.

Golden gate cloning was chosen for its capacity to efficiently generate diverse constructs using Type IIS restriction enzymes, which create non-palindromic, customizable overhangs. This feature is highly advantageous for the assembly of single-molecule constructs that require the attachment of distinct molecular handles at each terminus. The resulting versatility supports both plasmid- and linear-DNA experimental formats, enabling tailored construct design for specific applications. For example, in magnetic tweezers assays, asymmetric handles such as biotin and digoxigenin can be introduced at opposite ends, whereas symmetric overhangs restrict such modularity when using one restriction enzyme. When identical handles were required, fluorescent labeling on one or both sides can confer directionality and sequence identification (Figure 4B).

The protocol supported the successful synthesis of 5–100 kb constructs from donor plasmids, building on established methodologies (14,26,61), with improvements to accommodate the challenges posed by large DNA assemblies. Notably, using plasmid DNA as the starting material allowed for RecBCD pretreatment, which selectively removed linearized (damaged) DNA prior to digestion (Materials and Methods, Figure 3A). Subsequent size-exclusion chromatography eliminated excess handles, providing highly pure samples for fluorescence-based assays. Hybridized oligonucleotides with 5’ phosphorylation, biotin, or other modifications were used to introduce handles. Careful handling, with minimal pipetting and avoidance of vortexing, preserved DNA integrity during transition to AKTA purification. This approach consistently yielded high-quality and intact DNA, as verified by gel electrophoresis and fluorescence imaging (Figure 3B and C).

**Figure 3.**
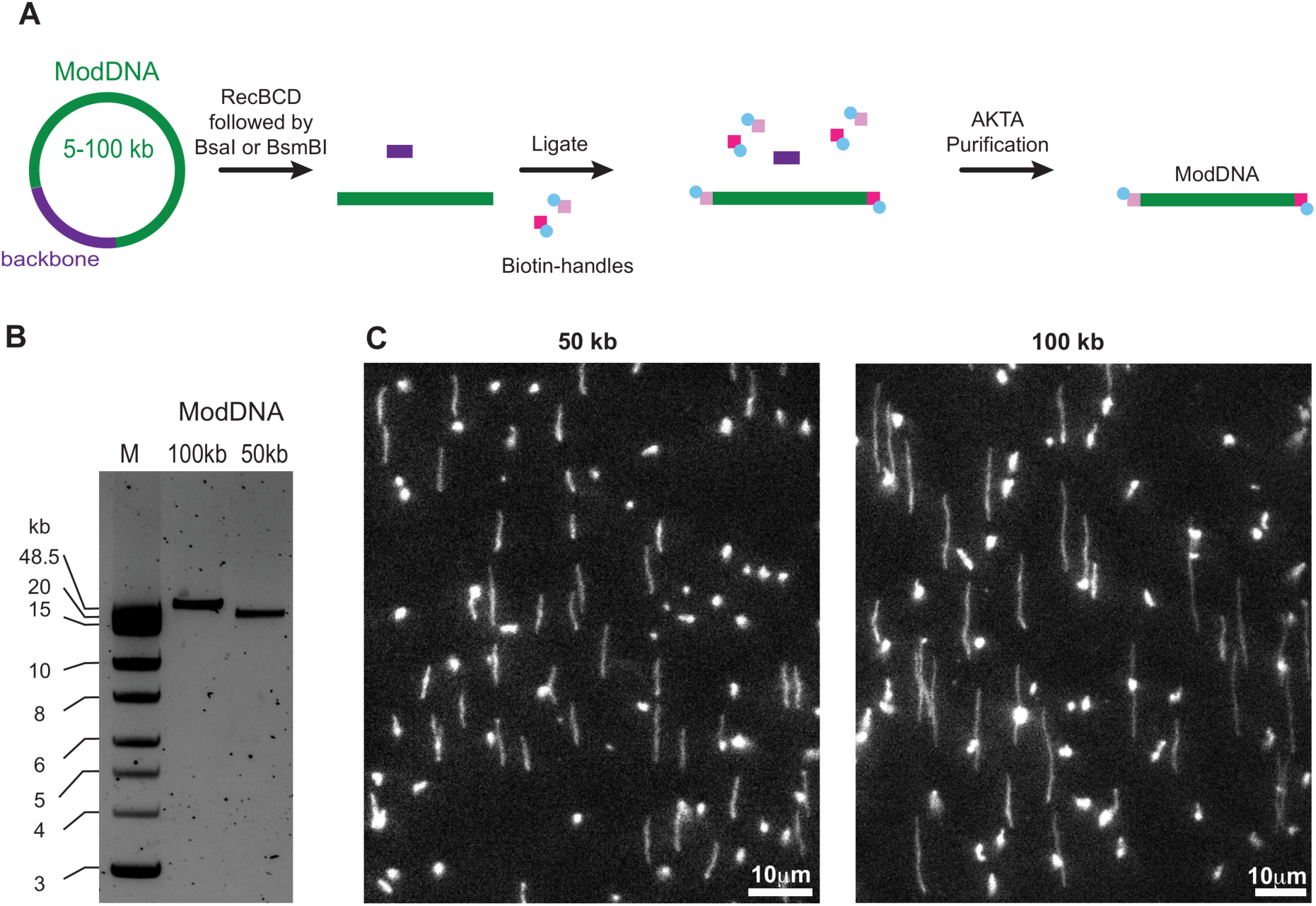
Production, purification, and single-molecule visualization of large ModDNA constructs. (A) Workflow for converting plasmid DNA into purified ModDNA constructs. To remove broken plasmids or contaminating genomic DNA, plasmids are first treated with RecBCD exonuclease, followed by digestion with Type IIS restriction enzymes, and ligation of additional biotin-handle DNA at the DNA ends. Non-ligated handles are subsequently removed by size-exclusion purification using an S500-HR column. (B) Finished and purified 50 kb and 100 kb ModDNA constructs were resolved on an agarose gel, yielding clean, well-defined DNA bands. (C) Fluorescence images of stretched, biotin-labeled ModDNA molecules, confirming the high purity of the ModDNA and suitability for single-molecule assays.

### Integration of local sequence features into ModDNA and single-molecule verification

In addition to enabling the creation of new custom-designed sequences, the ModDNA toolkit provides greatly enhanced flexibility over commonly used λ-DNA, permitting modification at any position within the sequence. Custom DNA sequences were introduced into the blank-slate ModDNA constructs by cloning them first into 5 kb Don2 vectors using standard techniques such as Golden Gate assembly, mutagenesis, or Gibson assembly (Figure 4A and Materials and Methods). Below, we present representative applications where we describe the engineered features (Figure 4) and single-molecule verification of them (Figure 4 and 5).

**Figure 4.**
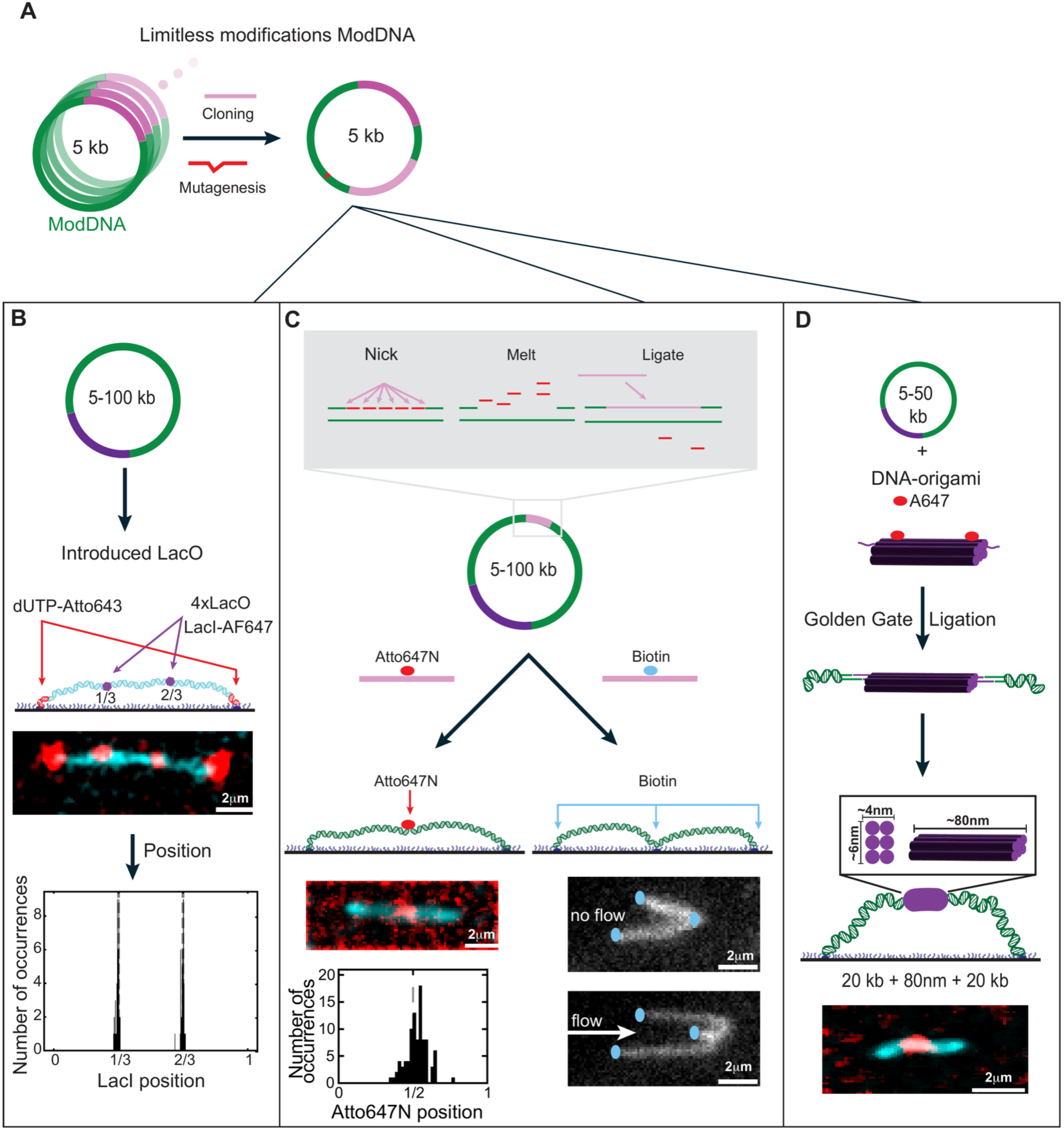
Functional modifications and engineered sequence features enabled by ModDNA. (A) Site-specific cloning enables targeted modifications of ModDNA. The subsequent panels show single-molecule example images of modular ModDNA constructs incorporating diverse functional elements. (B) A 50-kb ModDNA construct containing two 4×LacO sites positioned at 1/3 and 2/3 of its length, that exhibit specific and efficient LacI binding. (C) Incorporation of chemical and structural modifications, including mutagenized regions, chemical adducts, ssDNA or damage sites, and protein/peptide conjugates. Two examples are shown: (left) an Atto647N-labelled oligonucleotide ligated into the central gap of a 40 kb construct, with the distribution of measured label positions plotted below, and (right) insertion of an internal biotin at the same position, which tethers the molecule to the surface at a third point. (D) Using Type IIS restriction enzymes, an ∼80-nm DNA-origami structure can be ligated into the center of two 20-kb ModDNA fragments.

**Figure 5.**
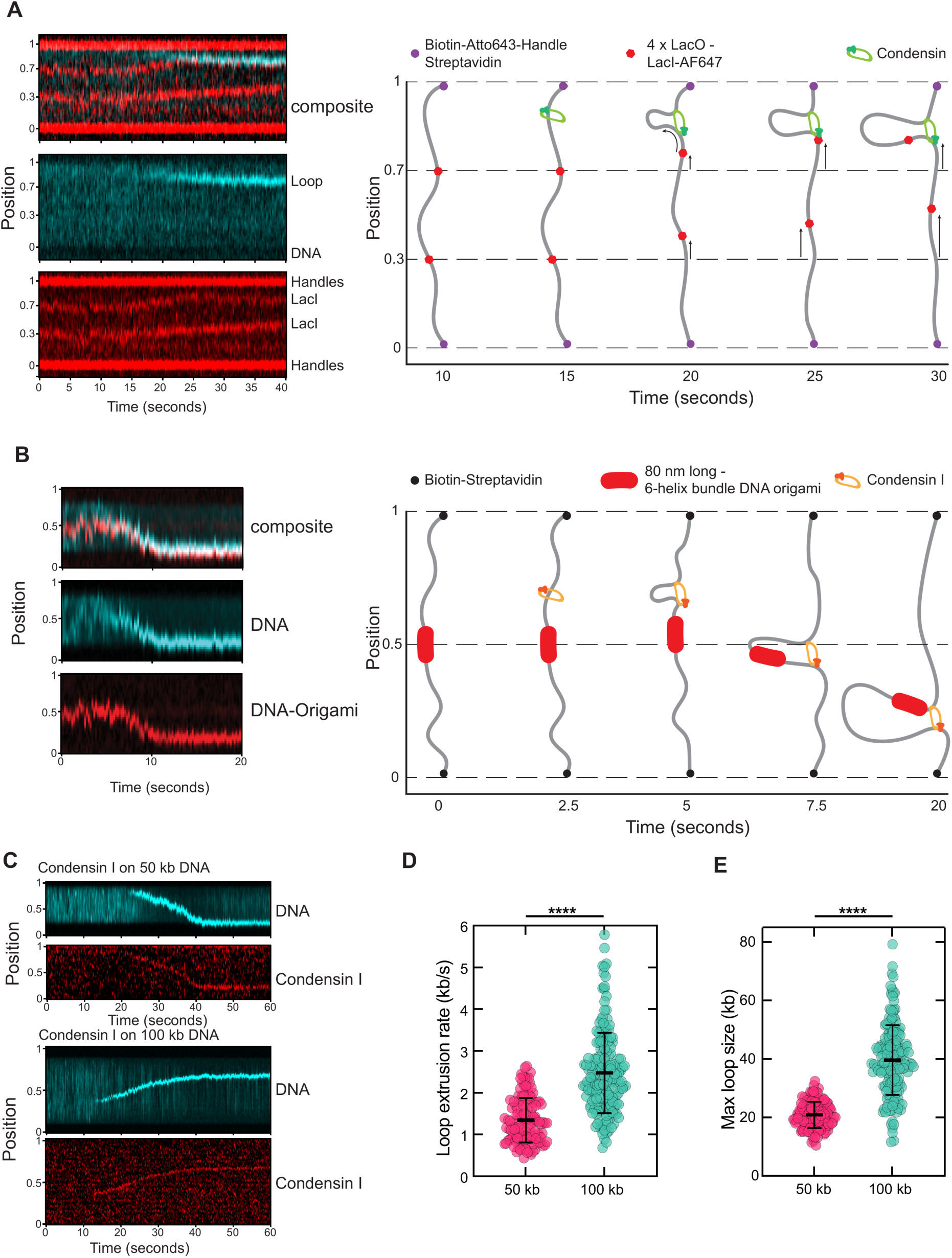
DNA loop extrusion by condensin on engineered ModDNA substrates. (A) Loop extrusion by budding yeast Condensin on ModDNA constructs carrying LacO/LacI labels at 1/3 and 2/3 positions (LacI-AF647; red), in the presence of SYBR Gold to visualize the DNA (cyan). The Atto643-labelled handles at the DNA ends are red in the kymographs and purple in the schematics. Left images show the experimental kymograph; right sketches illustrate the underlying molecular dynamics. Upon loop extrusion, the LacO/LacI labels move towards the loop. At a certain moment the LacO/LacI closest to the loop gets incorporated into the loop, whereupon it no longer moves in the kymograph, while the lower spot still moves up, indicating that the loop is still growing. (B) Loop extrusion by condensin I across an 80-nm DNA-origami barrier, with the DNA shown in cyan and the Alexa Fluor 647-labelled origami in red in both the kymographs and the schematics. The kymograph (left) shows a bypass event, with the schematics (right) illustrating the underlying molecular dynamics. (C) Kymographs of loop extrusion by condensin I on 50 kb (top) and 100 kb (bottom) ModDNA; the DNA (SYBR Gold, cyan) and condensin I (SNAP-Alexa Fluor 647, red) channels are shown separately. (D) DNA loop extrusion rates by condensin I on 50 kb (1.4 ± 0.5 kb/s) and 100 kb (2.5 ± 1.0 kb/s) ModDNA substrates (n = 131 for 50 kb and n = 173 for 100 kb; more than five independent replicates). (E) Maximum DNA loop sizes formed by condensin I on 50 kb (21 ± 5 kb) and 100 kb (40 ± 12 kb) substrates. In (D) and (E), bars represent the mean ± standard deviation. Statistical significance was assessed using the Mann-Whitney U test; both comparisons gave P < 0.0001, with effect sizes r = 0.72 and r = 0.86 for (D) and (E), respectively.

#### Site-specific fluorescent labelling and protein visualization

Two 4×LacO repeats were integrated at positions one-third and two-thirds along the length of a 50 kb ModDNA construct (Materials and Methods). We incubated this construct with 6 nM LacI-AF647, before immobilizing the DNA ends onto glass coverslips. Subsequently non-bound LacI-AF647 was washed out (Materials and Methods). This produced DNA molecules with two distinct fluorescent foci at the expected positions (Figure 4B), confirming that protein-binding sites can be placed at arbitrary, user-defined positions along the construct. Adding PCR-generated fluorescent handles labelled with Atto643–X–dUTP yielded constructs displaying four distinguishable spots, two at the ends and two at the defined LacO sites (Materials and Methods; Figure 4B).

This ModDNA construct can be used to monitor SMC-driven DNA loop extrusion in a dye-free manner, by tracking the position of the two LacI-bound spots (Figure 5A and S5A; Analikwu *et al.*, *in preparation*). Conceptually, this approach parallels dual-spot labelling that was recently used to observe cohesin-mediated loop extrusion in living human cells (62). DNA loop extrusion by SMCs *in vitro* is typically demonstrated and analyzed using intercalating dyes like SYTOX Orange, SYTOX Green, and others (12,13,16) but such dyes can alter the mechanics of DNA and quantitatively affect loop extrusion parameters (Analikwu *et al.*, *in preparation*). The same strategy, implemented here with LacO sites as a proof of principle, is broadly suitable for studies of enhancer–promoter interactions, transcription factors, and other DNA-binding proteins.

#### Nicking-enzyme cassettes for chemical handles and roadblocks

To place chemical modifications at defined internal positions, we adapted the nicking-and-replace strategy of Luzzietti *et al.* (54). Nb.BbvCI specifically cut one strand, where a series of recognition sites placed in the same orientation released the short intermediate ssDNA fragments, leaving a defined single-stranded gap. Using several closely spaced sites instead of a single distant pair meant that only short fragments had to dissociate, whereupon a long gap could be opened under mild conditions. An oligonucleotide spanning the whole gap, carrying any desired modification or mismatch, was then annealed and ligated, restoring an intact duplex that differs from the parent construct only at the intended position (Figure 4C, Materials and Methods).

Because BbvCI recognition sites were excluded from the ModDNA sequence during the in-silico design phase, we could introduce a BbvCI-cassette at any desired position. Here, we made a 40kb construct with five adjacent nicking sites at the center of the construct and used this construct to add three different modifications. First, an Atto647N containing oligonucleotide was used to assess the efficiency of integration. Of 132 double-tethered DNA molecules analysed, 94 carried the fluorophore (71.2%; 95% Wilson confidence interval 63.0-78.2%). Their measured label positions were centred on the expected position in the middle of the construct (Figure 4C). Instead of using an oligonucleotide carrying a pre-attached fluorophore, we tested the possibility of using an azide-containing oligonucleotide and labeled this with a DBCO-Alexa Fluor 647 (Jena Bioscience). As expected, the yield was lower, however, the measured yield is a combination of oligonucleotide integration, proportion of oligonucleotides with an azide, efficacy of azide-DBCO reactions and detection of Alexa 647 fluorophore which is less stable and bright than Atto647N. Here, 25 out of 69 molecules (36.2%; 95% Wilson confidence interval 25.9-48.0%) carried the signal. This illustrates both the feasibility of the click-based route and the need to verify directly that the intended modification is present on the DNA. Third, a biotinylated oligo was ligated into the middle gap. This produced DNA molecules that were attached to the streptavidin-coated surface at three spots: at both ends, but also in the middle. Imaging in lateral buffer flow confirmed the additional attachment of the middle position to the surface, confirming that the modification was correctly positioned and functional (Figure 4C, S5B). Because the replacement in this approach is an ordinary synthetic oligonucleotide, the same cassette accepts any modification that can be introduced during synthesis or attached through e.g. an azide. This includes peptides and protein conjugates, defined lesions or mismatches, and internal single-stranded regions.

#### Modular insertion of DNA origami

Type IIS restriction enzymes enabled modular integration of any object that can be ligated to DNA carrying appropriate 4-nucleotide overhangs. As a proof of concept, we inserted an 80 nm DNA origami 6-helix bundle, containing on either side a 4-nucleotide overhang, between two 20 kb ModDNA segments, which each already contained a biotin handle on one side (Materials and Methods, Figure 4D). Used as a roadblock in a loop-extrusion assay, the origami structure blocked condensin I in approximately 46 ± 20 % of events (95 % confidence interval; n=24; Figure 5B and S5C). Figure 5B shows an example where condensin I bypassed the DNA origami. Because the DNA origami was incorporated in-line, it forms a rigid segment of defined length within the DNA construct, mimicking an extended protein array covering a stretch of DNA. While SMC complexes can traverse DNA-attached particles up to 200 nm in diameter with a probability of ∼50% (53), the 80 nm-long origami achieved a comparable blocking rate despite being 2.5 times smaller. For arrays of telomeric Rap1 proteins bound to DNA in a similar manner as the DNA origami here, the blocking efficiency depended on the array length and on the size of the DNA gaps between bound proteins (18). By comparison, the origami roadblock is a continuous element with no gaps and a defined length. Given the modest 80 nm footprint of the bundle used here, a partial rather than complete blocking was to be expected. The same approach can accommodate pre-assembled nucleosome arrays, single-stranded fragments, or any other module carrying orthogonal overhangs.

#### Long DNA substrates for loop extrusion by SMC proteins

Next, we imaged loop extrusion by condensin I on the 50 kb and 100 kb constructs (Figure 5C and S6). DNA loop extrusion had so far not been studied *in vitro* on substrates of this length. Clear single-molecule loop extrusion events were observed on both, as seen in the kymographs of Fig. 5C. We measured a rate of 1.4 ± 0.5 kb/s (mean ± SD; N=131) on our 50 kb substrate (Figure 5D and S6A), similar to a 1.2 – 1.5 kb/s rate that was reported for condensin I on λ-DNA substrate (63). On the 100 kb substrates, we however measured a ∼1.8 fold higher rate than on 50 kb, 2.5 ± 1.0 kb/s (mean ± SD; N=173) (Figure 5D and S6C), starting from the same initial relative extension of the DNA (i.e. same initial tension in the DNA). Thus, the loop extrusion rate, surprisingly, appeared to be substrate-length dependent, a finding that we plan to further investigate with experiments and modelling in an independent study.

We did anticipate the loop size to be larger on 100 kb DNA than on 50 kb DNA. We therefore measured the maximum loop size that was reached during DNA loop extrusion on the 50 kb and 100 kb substrates. We found a maximum loop size that was about 1.9 fold larger on the 100 kb substrate than on the 50 kb construct, namely 40 kb ± 12 kb versus 21 ± 5 kb (Figure 5E and S6B and D). Both the loop extrusion rate and the maximum loop size were thus close to two-fold larger on the 100 kb substrate. The larger maximum loop size can be explained from the force build up on the double tethered DNA as illustrated by simulation curves (Figure S7). In these simulations, both the 50 kb and 100 kb were held at the same initial relative extension and hence loop extrusion started on them at the same low force (see also Materials and Methods, Figure S7). For every kilobase of DNA that was reeled in as loop extrusion proceeded, however, the relative extension of the DNA outside the loop rose twice as fast on the 50 kb substrate, and accordingly the internal force rose faster. Accordingly, the internal force increased to the level of the stalling force faster which terminated the process of loop extrusion. Or phrased otherwise, a loop can grow twice as large on the 100 kb construct to reach the same tension in the tether compared to a loop on the 50 kb construct.

## IV. Discussion and outlook

By developing a computational algorithm capable of generating random sequences that are depleted of any unwanted motifs (a “blank slate” DNA template) in highly modular long DNA, as well as by demonstrating a Golden Gate assembly workflow, we established a versatile new toolkit that has great potential for dissecting processes such as transcription, replication, and DNA damage repair on a wide variety of chromatin features such as CTCF sites, CpG islands, Widom 601 nucleosome arrays, enhancer–promoter elements, protein arrays, etc. Type IIS assembly and nicking-enzyme cassettes allow the inclusion or exclusion of elements such as binding sites, regulatory elements, defined lesions, gaps, replication-fork structures, etc., which all can be built into the substrate at chosen positions. Because the assembly is intrinsically modular, elements can easily be positioned anywhere along the ModDNA constructs, in any defined position and number. ModDNA thereby amply addresses the constraints that have limited λ-DNA as a single-molecule substrate (a fixed sequence, non-uniform GC distribution, a finite length, and little scope for internal modifications). Indeed, ModDNA can reach lengths (100 kbp demonstrated here, but potentially as long as ∼300 kbp) that were previously unattainable, while permitting full modularity and the ability to modify its sequence anywhere at will.

We anticipate that the increased length of the 100 kb substrate makes many single-molecule experiments much more tractable, for example studying observe collisions between SMC complexes, Z-loop formation (64) and other multi-complex experiments. The ModDNA system also has the added value that it can be combined with the modifications described above such as roadblocks, nucleosome arrays, or regulatory elements at defined positions along the DNA. ModDNA thus enables in-depth context-dependent dissection of SMC motor action at the single-molecule level *in vitro*. In some applications, one might like to focus on a limited set of criteria to build a DNA sequence that is tailored to address the research question at hand. For example, to build a prokaryotic-based synthetic genome, our lab is creating a sequence that is fully optimized for transcription by Escherichia coli RNA polymerase (H.H.A. Schenkel, in preparation). For this sequence, we do not limit the available sequence space by removing DNA motifs of yeast or human. Instead, we are particularly stringent in designing a ModDNA sequence that is free of *E. coli* features and additionally tailored in its DNA mechanical properties. We are planning to take the same approach for our studies with human SMC molecules, tailoring a sequence specifically to our research questions.

No doubt, databases, and the tools used to detect functional binding sites, will continue to evolve. Because of the modularity of ModDNA, iterations for new sequences can proceed through small mutations or by swapping out a single 5 kb region. The ModDNA pipeline was built for this. Its criteria can easily be sharpened and the synthesis is modular. The databases used here represent a defined snapshot, and updating a criterion requires only rerunning the design. The sequence reported here is in that sense the first member of a family of substrates rather than an ideal final sequence for random motif-free DNA.

The ability to generate long, motif-depleted DNA with fully defined sequence content will likely find applications beyond single-molecule assays. Indeed, ModDNA connects to three fast-moving scientific fields. First, the motif-aware design we use to *remove* unwanted elements is the mirror image of the (often machine-learning) approaches that are increasingly used to *design* functional regulatory DNA or promoters and enhancers with programmed activity built from learned sequence grammar (65–67). A curated, motif-depleted backbone then offers a defined “null” context in which such designed elements (and the polymerases and proteins that read them) can be characterized without confounding cryptic signals. Because the pipeline is organism-agnostic, the same algorithm can be retargeted to the regulatory logic of various species, for example, neutralizing or tuning promoters for the RNA polymerases of say yeast, mouse, or *E. coli*. The resulting sequences can be tested across scales, from bulk plate-reader expression assays to the single-molecule visualizations demonstrated here.

Second, motif-depleted sequences are attractive as neutral spacer or insulator elements in cell biology experiments, where long stretches of DNA must be included without introducing spurious transcription start sites, ribosome-binding sites, or open reading frames, which are precisely the cryptic elements that genome-scale design-and-synthesis efforts have to remove computationally (68–70).

Third, in a broad field called synthetic genomics, entire chromosomes are reengineered with systematic changes to coding sequences, regulatory elements, repetitive DNA, and genome architecture (71). The ModDNA capability to generate ‘clean’ DNA will no doubt be useful as a helpful tool for the re-engineering of more complex synthetic genomes, including emerging efforts to generate synthetic genomes for synthetic cells that are constructed from the bottom up (72). A validated route to long, programmable, motif-depleted DNA at the 100 kb scale is therefore directly relevant to synthetic-chromosome, synthetic-genome, and synthetic-cell engineering, in addition to its immediate use in single-molecule biophysics.

Overall, the ModDNA toolkit establishes a rigorous and adaptable foundation for long-DNA single-molecule assays, providing programmable sequence contexts for investigating protein-DNA interactions and the assembly of modular structural elements. The scalability and motif-aware design of ModDNA address the longstanding constraints of λ-DNA, while also complementing emerging synthetic biology assembly platforms. This adaptability positions ModDNA as a valuable resource for advancing a broad range of fundamental and applied studies in single-molecule biophysics and synthetic biology.

## Author Contributions

Conceptualization: JvdT, RB, BA, CD

Methodology: JvdT, RB, KM, CR, CD

Software: RB, BA

Experiments: JvdT, KM, TS, BA, CR, EB

Data analysis: JvdT, RB, TS, BA

Writing: JvdT, RB, KM, TS, BA, EB, BR, CD

Supervision: JvdT, BR, CD

## Acknowledgments

We thank Hilke Schenkel for discussions; Wouter De Laat for discussions and input on databases; Noa den Hartog, Amber van Landschoot, Jennate Elouach, Myrna de Jong, Lisanne Boere, Naomi Pijl, Kayleigh van Dongen, and Loïs van Gelder for contributions on testing, constructing, and cloning of ModDNA plasmids and constructs; Lisa Warners for her work on single-molecule DNA-Origami experiments and Victor Gilin for DNA-origami EM imaging. The pBeloBac11 plasmid was a kind gift from the lab of John van der Oost. We acknowledge the use of AI-based tools during manuscript preparation: Grammarly and Claude (Anthropic) for refining the text, and Claude for assistance in writing and checking the Python code used for analysis and modeling. All AI-assisted output was reviewed and verified by the authors, who take full responsibility for the content of this work.

