## Supplementary Information for "A modular DNA toolkit for single-molecule biophysics"

### Contents

This file contains one supplementary table and seven supplementary figures:

Table S1. Destination and donor vectors constructed in this study.

Figure S1. In silico sequence design: folding free energies, acceptance thresholds, and the resources screened.
Figure S2. Size-exclusion purification of DNA constructs on an S500-HR column.
Figure S3. Purification and characterisation of human condensin I.
Figure S4. Design and electron microscopy of the DNA-origami roadblock.
Figure S5. Additional single-molecule examples of engineered ModDNA constructs.
Figure S6. Quantification of loop extrusion by human condensin I on 50 kb and 100 kb ModDNA.
Figure S7. Worm-like-chain model of the DNA tension generated by loop extrusion.

Table S1

| Vector Type | Code | Plasmid # | Vector Name | Golden Gate | Creates |
|---|---|---|---|---|---|
| Destination | Dest1 | JT458 | pUC57mini-Amp-C0-C1 | BsmBI | + dsDNA = Don2 |
| | | JT459 | pUC57mini-Amp-C1-C2 | | |
| | | JT460 | pUC57mini-Amp-C2-C3 | | |
| | | JT461 | pUC57mini-Amp-C3-C4 | | |
| | | JT462 | pUC57mini-Amp-C4-C5 | | |
| | | JT463 | pUC57mini-Amp-C5-C6 | | |
| | | JT464 | pUC57mini-Amp-C6-C7 | | |
| | | JT465 | pUC57mini-Amp-C7-C8 | | |
| | | JT466 | pUC57mini-Amp-C8-C9 | | |
| | | JT467 | pUC57mini-Amp-C9-C10 | | |
| | | JT413 | pUC57mini-Amp-C10-C11 | | |
| | | JT414 | pUC57mini-Amp-C11-C12 | | |
| | | JT415 | pUC57mini-Amp-C12-C13 | | |
| | | JT416 | pUC57mini-Amp-C13-C14 | | |
| | | JT417 | pUC57mini-Amp-C14-C15 | | |
| | | JT418 | pUC57mini-Amp-C15-C16 | | |
| | | JT419 | pUC57mini-Amp-C16-C17 | | |
| | | JT420 | pUC57mini-Amp-C17-C18 | | |
| | | JT421 | pUC57mini-Amp-C18-C19 | | |
| | | JT422 | pUC57mini-Amp-C19-C20 | | |
| Donor | Don2 | JT246 | pUC57mini-Amp-C0-ModDNA-C1 | BsaI | + Dest2 = Don3<br>+ Dest3a = Don4a |
| | | JT247 | pUC57mini-Amp-C1-ModDNA-C2 | | |
| | | JT248 | pUC57mini-Amp-C2-ModDNA-C3 | | |
| | | JT249 | pUC57mini-Amp-C3-ModDNA-C4 | | |
| | | JT250 | pUC57mini-Amp-C4-ModDNA-C5 | | |
| | | JT251 | pUC57mini-Kan-C5-ModDNA-C6 | | |
| | | JT252 | pUC57mini-Amp-C6-ModDNA-C7 | | |
| | | JT253 | pUC57mini-Amp-C7-ModDNA-C8 | | |
| | | JT254 | pUC57mini-Amp-C8-ModDNA-C9 | | |
| | | JT255 | pUC57mini-Amp-C9-ModDNA-C10 | | |
| | | JT256 | pUC57mini-Amp-C10-ModDNA-C11 | | |
| | | JT257 | pUC57mini-Amp-C11-ModDNA-C12 | | |
| | | JT258 | pUC57mini-Amp-C12-ModDNA-C13 | | |
| | | JT259 | pUC57mini-Amp-C13-ModDNA-C14 | | |
| | | JT260 | pUC57mini-Amp-C14-ModDNA-C15 | | |
| | | JT261 | pUC57mini-Amp-C15-ModDNA-C16 | | |
| | | JT262 | pUC57mini-Amp-C16-ModDNA-C17 | | |
| | | JT263 | pUC57mini-Amp-C17-ModDNA-C18 | | |
| | | JT264 | pUC57mini-Amp-C18-ModDNA-C19 | | |
| | | JT265 | pUC57mini-Amp-C19-ModDNA-C20 | | |
| | | JT346 | pUC57mini-Amp-C4-5BbvCI-C5 | | |
| | | JT381 | pUC57mini-Amp-C3-4LacO-C4 | | |
| | | JT382 | pUC57mini-Amp-C6-4LacO-C7 | | |
| | | JT475 | pUC57mini-Amp-C11-ModDNA-C12_v2 | | |
| Destination | Dest2 | JT266 | pUC57mini-Kan-C0-C4 | BsaI | + Don2 = Don3 |
| | | JT312 | pUC57mini-Kan-C4-C8 | | |
| | | JT332 | pUC57mini-Kan-C8-C10 | | |
| | | JT359 | pUC57mini-Kan-C10-C12 | | |
| | | JT357 | pUC57mini-Kan-C8-C12 | | |
| | | JT341 | pUC57mini-Kan-C12-C16 | | |

| | | | | | |
|---|---|---|---|---|---|
| | | JT334 | pUC57mini-Kan-C16-C20 | | |
| Donor | Don3 | JT303 | pUC57mini-Kan-C0-ModDNA-C4 | BsmBI | + Dest3b = Don4b |
| | | JT333 | pUC57mini-Kan-C4-ModDNA-C8 | | |
| | | JT339 | pUC57mini-Kan-C8-ModDNA-C10 | | |
| | | JT361 | pUC57mini-Kan-C10-ModDNA-C12 | | |
| | | JT360 | pUC57mini-Kan-C8-ModDNA-C12 | | |
| | | JT342 | pUC57mini-Kan-C12-ModDNA-C16 | | |
| | | JT336 | pUC57mini-Kan-C16-ModDNA-C20 | | |
| | | JT385 | pUC57mini-Kan-C0-4LacO-C4 | | |
| | | JT386 | pUC57mini-Kan-C4-4LacO-C8 | | |
| Destination | Dest3a | JT314 | pBeloBacIlgga-C0-C8 | BsaI | + Don2 = Don4a |
| | | JT316 | pBeloBacIlgga-C0-C10 | | |
| | | JT323 | pBeloBacIlgga-C12-C20 | | |
| | | JT320 | pBeloBacIlgga-C0-C20 | | |
| Destination | Dest3b | JT315 | pBeloBacIlggb-C0-C8 | BsmBI | + Don3 = Don4b |
| | | JT317 | pBeloBacIlggb-C0-C10 | | |
| | | JT362 | pBeloBacIlggb-C10-C20 | | |
| | | JT321 | pBeloBacIlggb-C0-C20 | | |
| | | JT363 | pBeloBacIlggb-C12-C20 | | |
| Donor | Don4a | JT324 | pBeloBacIlgga-C0-ModDNA-C8 | BsmBI | + Dest3b = Don4b |
| | | JT351 | pBeloBacIlgga-C0-5xBbvCI-C8 | | |
| | | JT325 | pBeloBacIlgga-C0-ModDNA-C20 | | |
| Donor | Don4b | JT349 | pBeloBacIlggb-C0-ModDNA-C10 | BsaI | + Dest3a = Don4a |
| | | JT364 | pBeloBacIlggb-C10-ModDNA-C20 | | |
| | | JT365 | pBeloBacIlggb-C12-ModDNA-C20 | | |
| | | JT366 | pBeloBacIlggb-C0-ModDNA-C20 | | |
| | | JT387 | pBeloBacIlggb-C0-2x4LacO-C10 | | |

**Table S1. Destination and donor vectors constructed in this study.** For each vector, the table lists its class (destination or donor), its code in the ModDNA nomenclature (Dest1-Dest3b, Don2-Don4b), its plasmid number, its full name, the Type IIS enzyme used in the subsequent cloning step, and the product of that step. The enzyme used for Golden Gate assembly is the same enzyme used to release the insert when preparing a construct with biotin handles for single-molecule assays. Plasmid numbers correspond to the SnapGene files deposited with this work, which contain the complete sequence of every construct. The Don2 ModDNA donor vectors, with the exception of JT346, JT381, JT382 and JT475, and the Dest3 destination vectors were synthesised by GenScript; all remaining vectors were built in house by site-directed mutagenesis, Gibson assembly, or Golden Gate cloning. JT251 was supplied with a kanamycin rather than an ampicillin resistance cassette. This does not affect its use in the assembly and the marker was left unchanged. JT257 was functional but propagated poorly; JT475 carried a newly generated sequence for the same position (fr44 to fr47_v2), made with the same design pipeline so that it cloned and grew more easily, and it was not used for the data presented here.

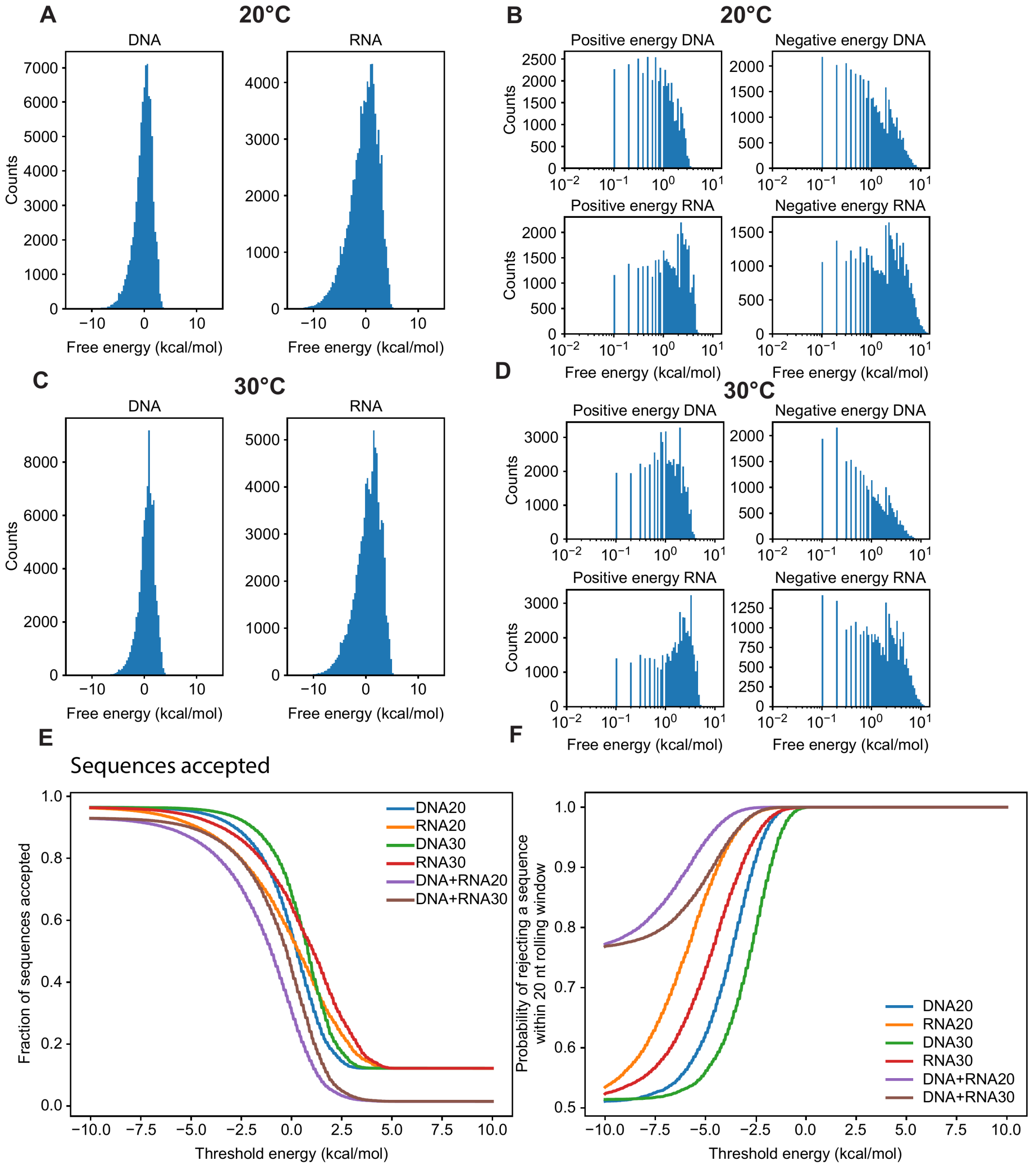


**G** Databases

| Databases | Source records |
|---|---|
| Promoters | 333,491 |
| Enhancers | 621,973 |
| Terminators | 5,251,075 |
| CollecTF | 5,854 |
| DHS | 8,313,473 |
| Curated RE | 5 |
| Repetitive | 4 |
| Total | 14,525,875 |

**H** Binding efficiencies, according to databases

| Databases | ModDNA | λ-DNA | Cosmid-Rap1 | Random |
|---|---|---|---|---|
| Sequence length (bp) | 100,000 | 48,502 | 41,965 | 100,000 |
| Promoters | 0 | 155 | 154 | 0 |
| Enhancers | 0 | 7 | 6 | 5 |
| Terminators | 0 | 186 | 173 | 3 |
| CollecTF | 0 | 314 | 277 | 592 |
| DHS | 10 | 15 | 22 | 22 |
| Curated RE | 0 | 27 | 22 | 111 |
| Repetitive | 2 | 2,076 | 1,770 | 3,052 |
| Total hits | 12 | 2,780 | 2,424 | 3,785 |

**Figure S1. In silico sequence design: folding free energies, acceptance thresholds, and the resources screened.** (A-D) Distributions of the predicted folding free energy of 100,000 randomly generated 20-nt sequences that had already passed all database-based exclusion criteria, computed with seqfold for both the DNA sequence and its RNA equivalent (Materials and Methods) at 20 °C (A, B) and 30 °C (C, D). Panels A/C show the full distributions on a linear energy axis; panels B/D show the positive- and negative-energy branches separately on a logarithmic axis. (E) Fraction of sequences accepted as a function of the rejection threshold, for DNA and for RNA at 20 °C and 30 °C, and for the combined DNA+RNA criterion at both temperatures. (F) Probability that a sequence is rejected within a 20-nt rolling window, for the same six criteria and as a function of the same threshold. Together, the results from panel E and F motivate the working threshold of $\Delta G < -3$ kcal/mol used throughout the design.

(G) Resources screened during sequence design: the number of source records taken from each database (Materials and Methods). (H) Sequence features detected in the finished ModDNA construct and in the three comparison sequences (λ-DNA, the Cosmid-Rap1 construct, and a random 100 kb sequence), given as the raw number of hits per database and in total (not corrected for sequence length, which is listed in the figure).

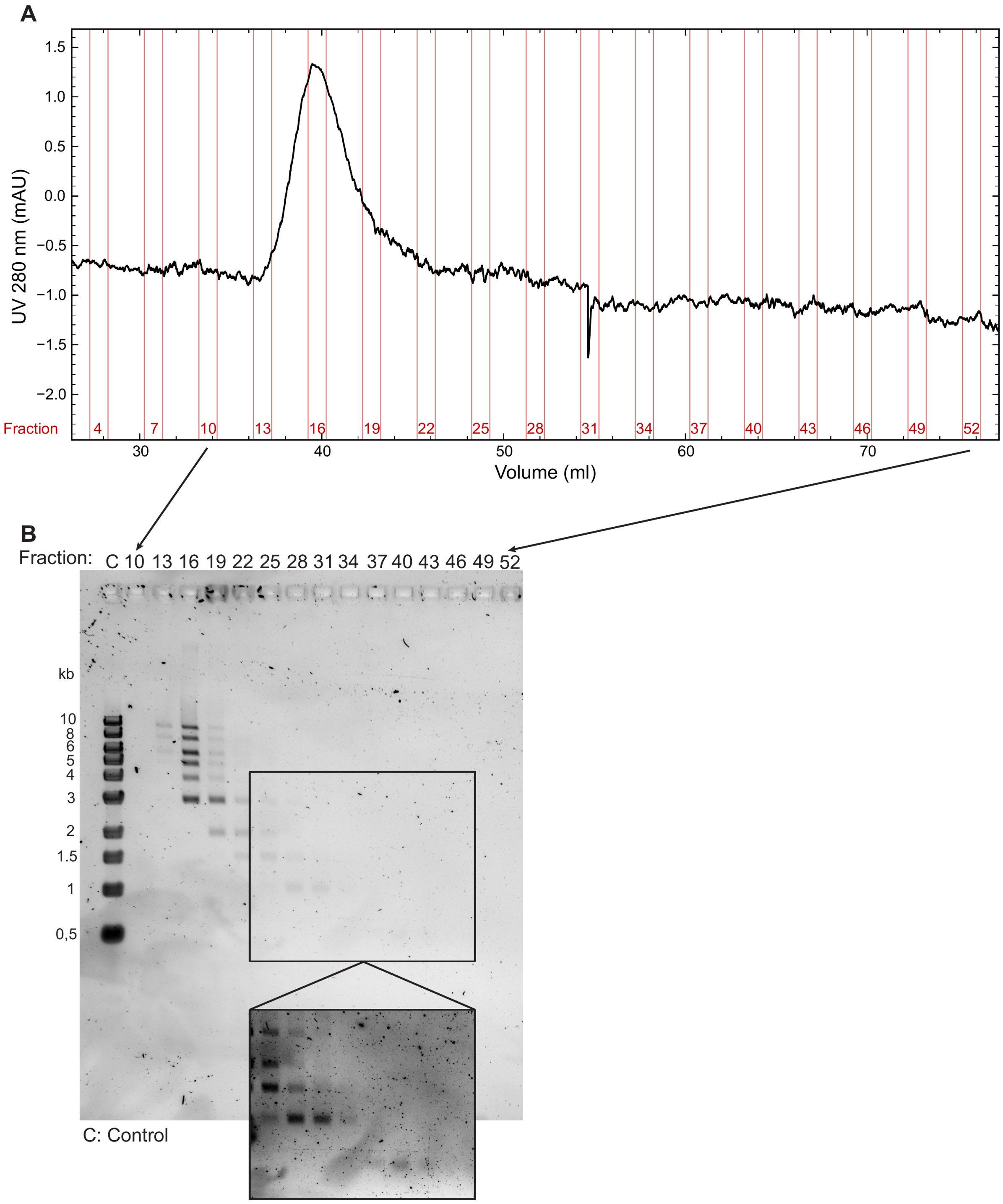


**Figure S2. Size-exclusion purification of DNA constructs on an S500-HR column.**

(A) UV absorbance at 280 nm during a representative AKTA run of a 1 kb DNA ladder (NEB) on an S500-HR column (Cytiva) in TE supplemented with 150 mM NaCl at 0.5 ml/min. Vertical lines mark some of the collected fractions. (B) Agarose gel of collected fractions. The high-molecular-weight species elute first and progressively smaller fragments elute later, so that free handles and unligated oligonucleotides are separated from the assembled construct. The boxed region is shown enlarged and at higher contrast at the bottom of the gel. Lane C provides a control (input ladder).

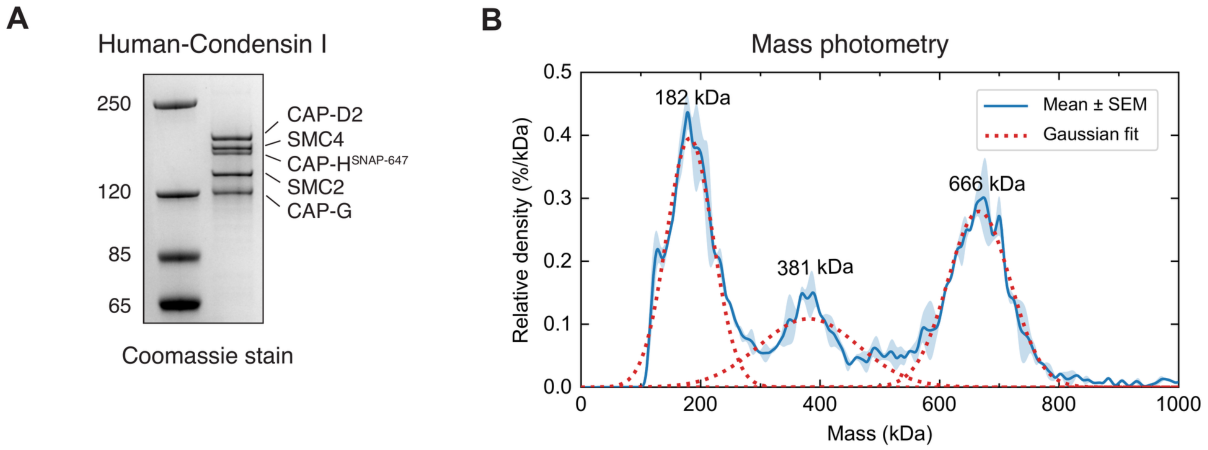


**Figure S3. Purification and characterisation of human condensin I.**

(A) Coomassie-stained SDS-PAGE gel of the human condensin I preparation, with the subunits indicated (CAP-D2, SMC4, CAP-H-SNAP-647, SMC2, CAP-G). (B) Mass photometry of the human condensin I preparation (blue, mean ± SEM of n = 2 measurements; red dotted line denotes a Gaussian fit). The peak at 666 kDa corresponds to the complete SNAP-tagged holocomplex (calculated mass 656 kDa); the peaks at 381 and 182 kDa correspond to noncomplete holocomplexes.

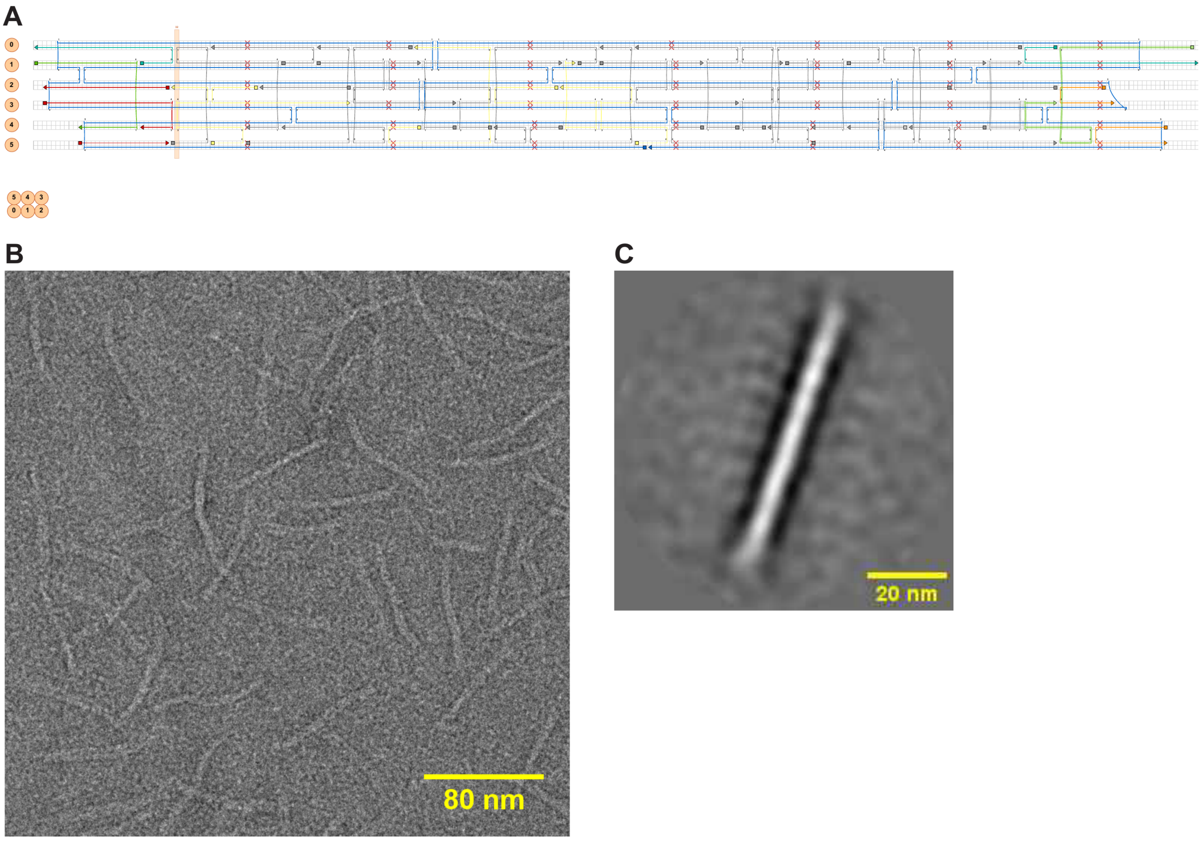


**Figure S4. Design and electron microscopy of the DNA-origami roadblock.**

(A) caDNAno design of a 3x2-helix bundle that is used as the roadblock in SMC-driven DNA loop extrusion. The structure is folded on a 1426-nucleotide scaffold excised from p7560 by MscI and HincII digestion, with a skip inserted every 32 nucleotides in each helix to compensate for twist (Materials and Methods). (B) Transmission electron micrograph of the purified DNA origami; scale bar 80 nm. (C) Averaged image of the folded structure, showing the expected rod-like bundle of ~80 nm length. Scale bar 20 nm.

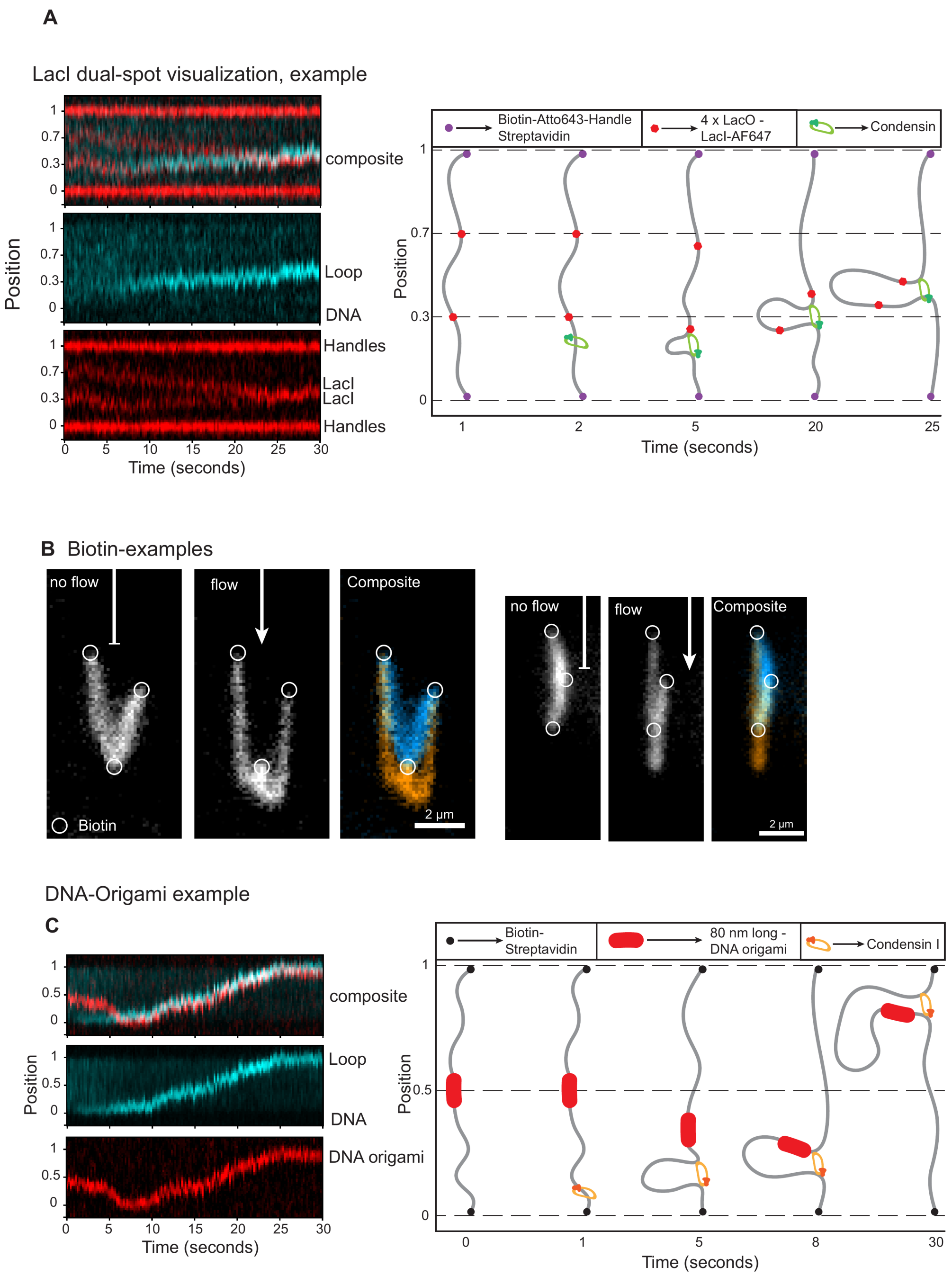


**Figure S5. Additional single-molecule examples of engineered ModDNA constructs.**

(A) Dye-free tracking of loop extrusion on a 50 kb ModDNA construct carrying two 4xLacO arrays, one at one third and one at two thirds of its length. Left, kymographs of the composite, showing the DNA channel (SYBR Gold) and the red channel that reports both the biotinylated Atto643 end handles and the Alexa647-labelled LacI bound to the LacO arrays. Right, schematic of the DNA at the indicated times, showing that the LacI-bound arrays move as extrusion proceeds. (B) Examples of a 40 kb ModDNA construct that carries an internal biotin, imaged without and with buffer flow. Circles indicate the three points at which the molecule is attached to the streptavidin-coated surface. (C) Loop extrusion by human condensin I on a 20 kb-origami-20 kb construct. Left, kymographs; right, schematic at the indicated times.

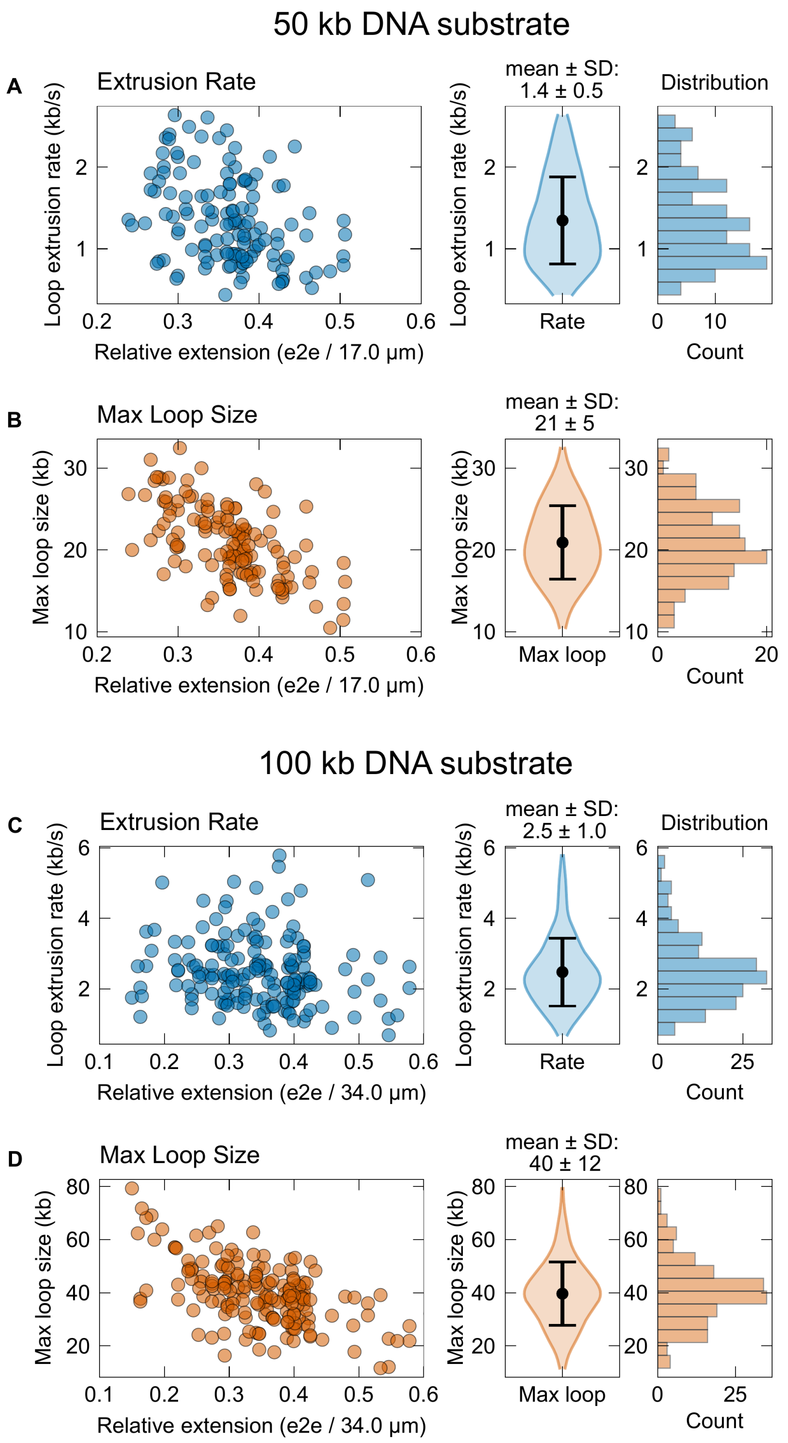


**Figure S6. Quantification of loop extrusion by human condensin I on 50 kb and 100 kb ModDNA.** (A, C) Loop extrusion rate as a function of the initial relative extension of the DNA molecule (end-to-end distance divided by the contour length, which is 17.0 µm for the 50 kb and 34.0 µm for the 100 kb substrate), on a 50 kb (A) and 100 kb (C) substrate. Mean ± SD was 1.4 ± 0.5 kb/s and 21 ± 5 kb on the 50 kb substrate (n = 131), and 2.5 ± 1.0 kb/s and 40 ± 12 kb on the 100 kb substrate (n = 173), from more than five independent replicates. (B, D) The same representation for the maximum loop size reached during the trace, on the 50 kb (B) and 100 kb (D) substrates.

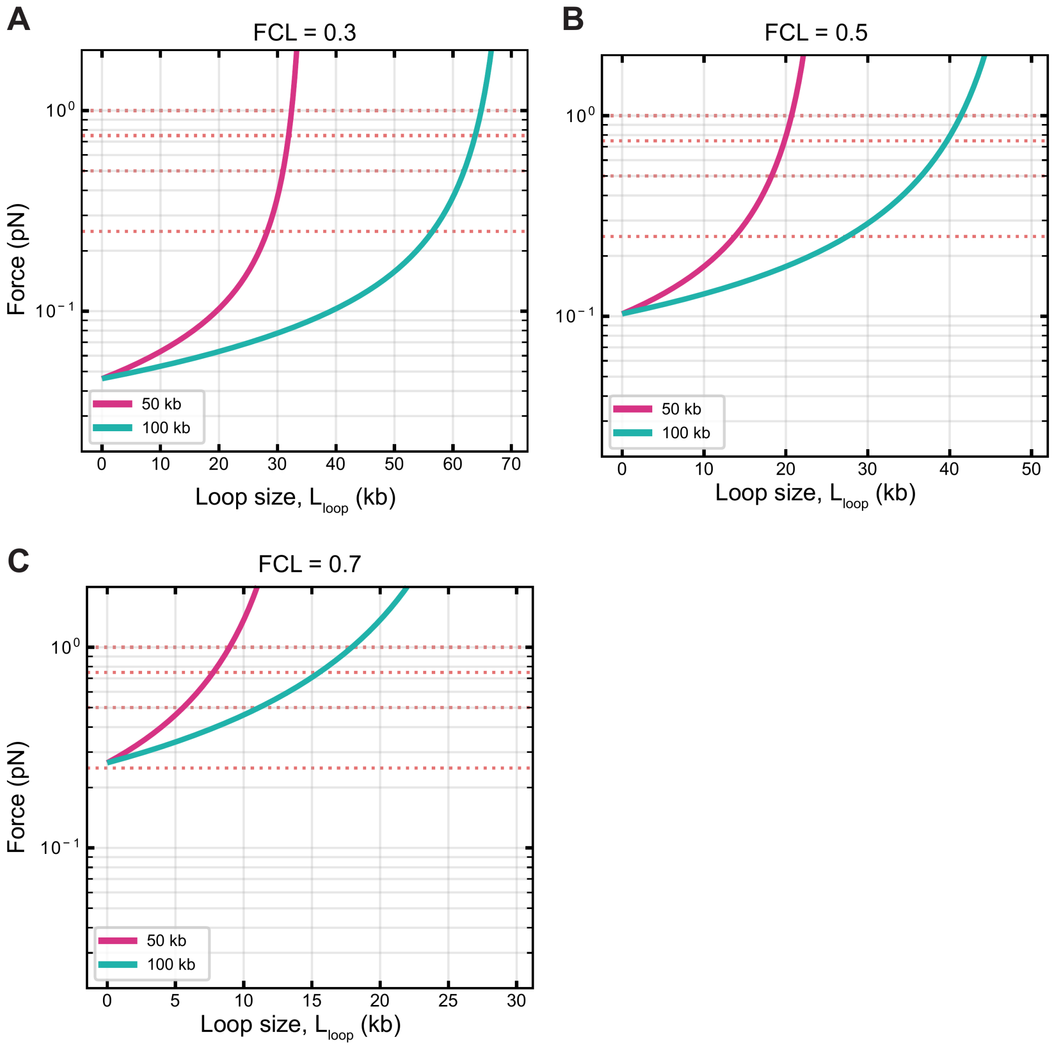


**Figure S7. Worm-like-chain model of the DNA tension generated by loop extrusion.**

Force in the DNA tether outside the loop as a function of extruded loop size, for 50 kb (magenta) and 100 kb (teal) molecules held at a fixed end-to-end distance, calculated from the Marko-Siggia interpolation formula (Materials and Methods). (A) FCL = 0.3, (B) FCL = 0.5 and (C) FCL = 0.7. Red dotted lines mark reference forces of 0.25, 0.5, 0.75 and 1 pN. Notably, at the end of the loop extrusion, the same tension is reached at twice the absolute loop size on the 100 kb substrate as compared to on the 50 kb substrate.
